# A Highly Conserved *Shh* Enhancer Coordinates Hypothalamic and Craniofacial Development

**DOI:** 10.1101/794198

**Authors:** Zoe Crane-Smith, Jeffrey Schoenebeck, Katy A Graham, Paul S Devenney, Lorraine Rose, Mark Ditzell, Eve Anderson, Joseph I Thomson, Natasha Klenin, Deborah M Kurrasch, Laura A Lettice, Robert E Hill

## Abstract

Enhancers that are conserved deep in evolutionary time regulate characteristics held in common across taxonomic classes. Here, deletion of the highly conserved *Shh* enhancer SBE2 (*Shh* brain enhancer 2) markedly reduced *Shh* expression within the embryonic brain specifically in the rostral diencephalon; however, no abnormal anatomical phenotype was observed due to compensatory low levels of expression mediated by a second enhancer. In contrast, a further reduction of *Shh* levels, achieved by crossing the SBE2 deletion with the *Shh* null allele, disrupted brain and craniofacial tissue; thus, linking SBE2 regulated *Shh* expression to multiple defects and further enabling the study of the effects of differing levels of *Shh* on embryogenesis. Development of the hypothalamus, derived from the rostral diencephalon, was disrupted along both the anterior-posterior (AP) and the dorsal-ventral (DV) axes. Expression of DV patterning genes and subsequent neuronal population induction were particularly sensitive to *Shh* expression levels, demonstrating a novel morphogenic context for *Shh*. The role of SBE2, which is highlighted by DV gene expression, is to step-up expression of *Shh* above the minimal activity of the second enhancer, ensuring the necessary levels of Shh in a regional-specific manner. We also show that low *Shh* levels in the diencephalon disrupted neighbouring craniofacial development, including mediolateral patterning of the bones along the cranial floor and viscerocranium. Thus, SBE2 contributes to hypothalamic morphogenesis and ensures there is coordination with the formation of the adjacent midline cranial bones that subsequently protects the neural tissue.

## Introduction

Long-distance enhancers that are conserved across multiple classes of vertebrates are implicated in regulating early embryonic development (Nord et al., 2013). The *Shh* regulatory domain, which extends 1Mb upstream of the gene contains a number of sequence elements identified by high sequence conservation. These elements appear to be largely responsible for the typical pattern of *Shh* expression observed at initial stages of organogenesis (Anderson et al., 2014; Jeong et al., 2008). The deep conservation and early stages of expression mediated by these *Shh* enhancers suggest a fundamental role in generating ancient, shared vertebrate characteristics. Studies to date show a modular composition of the *Shh* regulatory domain and thus, for at least some of the *cis*-regulators, there is a simple relationship of a single enhancer to a distinct spatial domain of expression (Jeong et al., 2006; Lettice et al., 2014); although, in other instances, secondary enhancers provide compensatory low levels of expression (Letelier et al., 2018; Tsukiji et al., 2014).

SHH is a signalling factor with a fundamental role in brain and craniofacial development. A number of enhancers have been identified within the *Shh* regulatory domain that are responsible for regulating expression in the midbrain and forebrain. Transgenic analysis indicates that the SBE2 enhancer (Jeong et al., 2008, 2006; Zhao et al., 2012) regulates *Shh* expression in the early developing forebrain, from the zona limitans intrathalmica (ZLI) to the medial ganglionic eminence (MGE), specifically designated the rostral diencephalon (Jeong et al., 2006; Zhao et al., 2012). Of particular interest is a clinically relevant inactivating mutation in the SBE2 enhancer associated with features of semilobar holoprosencephaly (Jeong et al., 2008). Holoprosencephaly (HPE) is a developmental disorder of both the forebrain and midline facial structures that include microcephaly, midfacial hypoplasia, cleft lip and palate, diabetes insipidus and moderate fusion of the hypothalamus (Marcorelles and Laquerriere, 2010). More commonly, HPE is associated with heterozygous mutations that inactivate the *Shh* gene (Chiang et al., 1996; Ming and Muenke, 2002). These data suggest that disruption of SBE2 driven expression in the early developing forebrain may contribute to this disorder (Jeong et al., 2008).

Here, we deleted the SBE2 enhancer to discern its overall contribution to *Shh* expression pattern and to establish its requirement for mediating the embryonic phenotype. *Shh* expression is continuous throughout the central nervous system, relying on a number of *cis*-regulators to generate the overall pattern (Anderson et al., 2014). The deletion of SBE2 interrupted this spatial pattern by specifically downregulating expression in the ventral rostral diencephalon; however, this reduction of SBE2 driven *Shh* expression produced no overt phenotype. We confirmed the presence of a second enhancer (Sagai et al., 2019) responsible for residual levels of *Shh* expression that compensated for the loss of SBE2 in the rostral diencephalon. Further reduction in expression levels of *Shh* was attained by crossing the SBE2 deletion mice to the line carrying the *Shh* null allele. The development of the hypothalamus and of the surrounding tissue was disrupted in the compound mutants, linking diencephalic spatiotemporal expression to multiple defects and enabling analysis of development at different expression levels. We showed, firstly, that patterning in the hypothalamus along the AP axis was disrupted, but low levels of *Shh* are sufficient for normal formation of the AP boundary. Further reductions in the levels of *Shh*, however, led to misplaced spatial assignment of the hypophyseal lobes due to disruption of the AP boundary. Secondly, gene expression along the DV axis was, in contrast, concentration sensitive and consistent with SHH functioning as a morphogen along this developmental axis, with tissue patterning and eventual neuronal fate reliant upon varying concentrations of *Shh* signalling. Thirdly, we found that loss of SBE2 activity disrupts craniofacial development affecting medial lateral patterning of the bones along the cranial floor and palatal shelf, revealing that SBE2 activity in the brain directly regulates cranial and facial morphogenesis, providing further evidence of the ventral patterning role played by *Shh* (Corman et al., 2018; Dworkin et al., 2016). These data confirm that signalling in the brain has an influence upon craniofacial morphology and raises an interesting evolutionary relationship that links patterning of a crucial region of the brain with the formation of the bony plate barrier underlying this region in the skull. Hence, a single regulatory component coordinates the combined function of directing brain development with the concomitant role of ensuring physical neuroprotection.

## Materials and methods

### Embryo production and transgenic targeting

All animal studies were approved by the MRC IGMM ‘Animal Care and Use Committee’ and according to the MRC ‘Responsibility in the Use of Animals for Medical Research’ (July 1993), EU Directive 2010 and UK Home Office Project License no PPL 60/4424.

Embryos were harvested at various embryonic stages between E10.5-E17.5. For all experiments described triplicate datasets were used, unless otherwise stated. Reduction in *Shh* levels to 50%, in *Shh*^*null*/+^ embryos, is known to cause no phenotypic effects (Chiang et al., 1996). Nevertheless, to allow for the use of littermate controls, differences between wild-type, *Shh*^*ΔSBE2*/+^ and *Shh*^*null*/+^ embryos were assessed for each experiment described. As expected, no differences were detected in between the three previously mentioned genotypes, as such all three genotypes were used as control embryos.

The expression driven by potential enhancer region of interest was analysed in G0 transgenic embryos using a lacZ construct which carried sequence of the region in question, a β-globin minimal promoter and LacZ as has been previously described (Yee and Rigby, 1993) The DNA fragment of interest was generated using PCR with the primers GATCAT**GTCGAC**GCTCCAGGTACTGCTGTTCAG and GATCAT**GCTGAC**ATGTGGATGGCAAGCATTGGC- the SalI restriction site used in the cloning is highlighted in bold.

### ATAC-seq and FAC sorting

Entire heads of GFP positive eGFP/Cre-*Shh* (Harfe et al., 2004) embryos were dissected and single cell suspensions were made by incubating the dissected tissue in 1:5 dilution of trypsin:versene at 37°C for 15 minutes. GFP positive cells were sorted using a fluorescence activated cell sorter and ATAC-seq libraries were made from the positive cell population pooled across littermates as previously described (Buenrostro et al., 2015). Subsequently libraries were subject to size selection to exclude DNA fragments larger than 1 kb using SPRIselect magnetic beads (Beckman Coulter). Samples were sequenced on a Illumina HiSeq 4000 platform to obtain 50 bp paired-end reads. Resulting reads were trimmed using cutadapt paired-end trimming, and subsequently aligned to the mm9 mouse genome using bowtie2 paired-end alignment. Unmapped reads and those mapping to the mitochondrial genome were then removed and duplicate reads were filteres out using Picard MarkDuplicates. Reads were shifted by +4bp for those mapping to the positive strand and −5bp for those mapping to the negative strand. Broad peaks were then called, to identify open chromatin regions, using MACS2 callpeak, using –g mm, -f BEDPE, -B –broads and cutoff 0.01 options.

### SBE2 deletion line production by CRISPR

CRISPR guide oligo pairs were designed flanking the SBE2 enhancer region described by Zhao, *et al* 2012. The pairs were annealed and cloned into the pX330 vector. Guide sequences used were: Upstream of SBE2, 5’ AAACACATTAAAGCCCTCCAGCG 3’ and 5’ CACCGACGCTGGAGGGCTTTAATGT 3’; Downstream of SBE2, 5’ AAACACGAGCAAGCCAACCGGAGG 3’ and 5’ CACCGCTCCGGTTGGCTTGCTCGT 3’. pX330-U6-Chimeric_BB-CBh-hSpCas9 was a gift from Feng Zhang (Addgene plasmid # 42230; http://n2t.net/addgene:42230; RRID:Addgene_42230) (Cong et al., 2013).

Transgenic mice were made by pronuclear injection of plasmid DNA into C57Bl/6 x CBA F2 embryos. G0 pups were genotyped by PCR and sequencing. Pups carrying the correct deletions were then used to establish the line by mating to C57BL/6J mice.

### OPT analysis

OPT imaging was performed (Sharpe, 2003). Briefly, PFA-fixed embryos were embedded in 1% low-melting-point agarose and dehydrated by immersion in methanol for 24 h. The sampes were cleared for 24 h in BABB (one part benzyl alcohol/two parts benzyl benzoate). before being scanned using a Bioptonics 3001 scanner (www.biooptonics.com) Images taken every 0.9° (of a 360° rotation) and were reconstructed using Bioptonics proprietary software with the outputs then being viewed with Dataviewer (Bioptonics) and Bioptonics Viewer.

### Morphometric analysis

MicroCTs of fifteen E17.5 embryos (7 controls, 8 mutants) collected from three litters were scanned post mortem using a Bruker Skyscanner 1076. Individual skull scans were downsampled 2:1 using Bruker NRecon. Reconstructed images were converted to DICOM format using Bruker DicomCT. Twenty-eight landmarks were placed on each isosurface using Stratovan CheckPoint (v19.03.04.1102 Win64). In two mutants, dysmorphology of the vomer prevented placement of corresponding landmarks and as such, the landmarks were indicated as “missing”. Raw coordinates data saved in Morphologika format were processed in R (v3.5.3) using Geomorph (v3.1.2), Rvcg (v0.18), Morpho (v2.7), and Hotelling (v1.0-5) packages. Missing landmarks were estimated using a multivariate regression method implemented in Geomorph. Following, a Generalised Procrustes analysis (GPA) translated, rotated and rescaled raw landmark coordinate data to minimise the sum of squared distances between specimens. The amount of rescaling required in the GPA is captured by the centroid size, a variable which is used subsequently as a proxy for specimen size. A single outlier (control, Litter 3) was removed as its Procrustes distance was >1 standard deviation from the mean. Following removal of the outlier, the GPA was rerun to produce Procrustes shape variables. To test for shape variation due to allometry, shape variables were regressed against log(centroid size) as well as additional terms corresponding to genotype and litter. The best fit included log(centroid size) interactions with both litter and genotype (ie coordinates ~ log(centroid size)*litter + log(centroid size)*genotype).

In order to visualise shape differences, a randomly chosen ply surface file was warped using thin plate splines using the mean coordinate values of all fourteen specimens. The resulting mesh represented the dataset reference which was subsequently warped to using principal component minimum and maximum shape variables.

#### 1.1.1 Tissue processing

Embryos were dissected at the stages indicated and fixed overnight at 4°C in 4% paraformaldehyde (PFA) in phosphate-buffered saline (PBS). Embryos were washed briefly in PBS, and then passed through a sucrose gradient: 30 min in 5% sucrose in PBS, 30 min in 10% sucrose in PBS before being placed in 20% sucrose in PBS overnight at 4°C. Embryos were embedded in OCT compound and stored at −80°C. Sections were cut on a cryostat at varying thicknesses appropriate to the embryonic stage: For E11.5 at 8 μm, for E13.5 at 10 μm and for E17.5 at 14 μm.

#### 1.1.2 H&E staining

Cryosections were allowed to reach room temperature before the slides were rehydrated by passage through 100% EtOH 3x 5 min then serial passage through 90%, 70%, 50% and lastly 30% EtOH for a couple of min each. Slides were washed in H_2_O and stained in haemotoxylin for 4 min. They were rinsed in H_2_O and subsequently differentiated in acid/alcohol for a few seconds. Slides were again rinsed in H_2_O then placed in saturated lithium carbonate solution for a few seconds. They were rinsed in H_2_O, stained in eosin for 2 min and again rinsed in H_2_O. Slides were dehydrated through 4 changes of 100% ethanol and passaged 3x through xylene for 5 min each, subsequently mounted in DPX (Cell Path, SEA-1304-00A), coverslipped and allowed to dry overnight.

### *lacZ* expression analysis, Dawson staining and wholemount *in situ* hybridisation

Embryos were analysed for *lacZ* expression at E10.5 through staining for β-gal activity as previously described (Mackenzie et al., 1997). Skeletal preparations from E17.5 foetuses were stained simultaneously with Alizarin Red and Alcian Blue (Nagy et al., 2009a, 2009b). Whole-mount *in situ* hybridisation on half heads was performed as previously described (Hecksher-Sørensen et al., 1998). Whole-mount *in situ* hybridisation on E95 day embryos was performed as previously described (Pai et al., 2015).

### *In situ* hybridisation on cryosections

For in situ on cryosections, the protocol was adaped from (Lu et al., 2013) as described. Briefly, *in situ* probes were placed at 37°C for 5 mins, added to hybridisation buffer (1x salt (2 M NaCl, 100 mM Tris, 65 mM NaH_2_PO_4_, 50 mM Na_2_HPO_4_ and 50 mM EDTA in ddH_2_O, pH to 7.5), 50% deionised formamide, 10% dextran sulfate, 1 mg/mL yeast RNA, 1x Denhardt’s (2% BSA, 2% Ficcoll, and 2% PVP made up to 100mL with ddH_2_O)) pre-warmed to 65°C and subsequently denatured at 95°C for 2 mins. Probes were added to slides, that had been previously to reach room temperature, and placed in a 65°C waterbath overnight in a chamber humidified with 5x SSC/50% formamide. Slides were subject to 3x 30 min washes at 65°C in washing solution (50% formamide, 25 mL 1x SSC and 0.1% Tween 20 made up to 500 mL Milli-Q H_2_O). Subsequently, slides were washed 2x 30 min in 1x MABT (100 mM maleic acid, 167 mM NaCl in ddH_2_O, pH to 7.5, then 0.1% Tween 20). Slides were blocked in 1x MABT + 2% blocking reagent + 20% heat inactivated sheep serum. A 1/2500 dilution of anti-DIG (Roche, 1109327490) in 1x MABT + 20% heat inactivated sheep serum was then added and slides were placed in a humified chamber and incubated overnight at room temperature. Slides were washed 5x for 20 min in 1x MABT. They were then washed with NTMT (100 mM NaCl, 50 mM MgCl, 100 mM Tris pH 9.5 and 0.1% Tween 20 made up in ddH_2_O) for 10 min followed by NTMTL (NTMT + 5 mM Levamisole) for 10 min. Staining solution, NTMTL containing 3.75 μL/mL BCIP (Roche, 11383221001) and 5 μL/mL NBT (Roche, 11383213001), was then added and slides were placed at 37°C to allow colour to develop. Slides were immersed in H_2_O, air dried and mounted in Prolong gold (Thermo Fisher, P39634).

#### 1.1.3 Immunofluorescence

Cryosections were removed from the −80°C freezer and allowed to reach room temperature before being subjected to antigen retrieval using a citric acid solution (10 mM citric acid dissolved in ddH_2_O, then pH to 6-6.5). Subsequently, slides were washed twice for 5 min in PBS then blocked using 10% goat serum in PBT (0.1% Triton-X100 in PBS) for 1 h at room temperature. For PITX2 staining, donkey serum was used, Slides were incubated in primary antibodies diluted in blocking solution overnight at 4°C as follows: rabbit anti-CASP3 (1:500, Abcam, ab49822); mouse anti-COL2A1 (1:400, Santa Cruz, sc-52659); rabbit anti-ISL1 (1:200, Abcam, ab20670); rabbit anti-LHX3 (1:250, Abcam, ab14555); rabbit anti-NKX2-1 (1:250, Abcam, ab76013); rabbit anti-NKX2-2 (1:250, Abcam, ab191077); rabbit anti-NKX6-1 (1:250, Abcam, ab221549); rabbit anti-OLIG2 (1:200, Abcam, ab109186); rabbit anti-oxytocin (1:500, Abcam, ab212193); rabbit anti-PAX6 (1:350, Abcam, ab195045); sheep anti-PITX2 (1:200, R&D, AF7388); rabbit anti-SF1 (1:200, Abcam, ab65815); rabbit anti-TBX3 (1:200, Abcam, ab99302); rabbit anti-TH1 (1:200, Abcam, ab137869); rabbit anti-TCF4 (1:100, Cell Signalling; C48H11); rabbit anti-vasopressin (1:500, Abcam, ab213707). Slides were then washed 3 times for 10 min each in PBT and incubated for 1 h at room temperature with fluorescent secondary antibodies in PBT in the dark, secondaries as follows: goat anti-rabbit 488 (1:500, Thermo Fisher, A11034); goat anti-mouse 488 (1:500, Thermo Fisher, A11029); donkey anti-sheep 594 (1:500, Thermo Fisher, A11016). Slides were then washed 3 times for 10 min each in PBT and subsequently stained with 0.1 μg/ml DAPI in PBS for 10 min, then washed in PBS twice for 5 min each. Lastly, slides were mounted using Prolong Gold (Thermo Fisher, P39634) and allowed to dry prior to imaging.

## RESULTS

### Loss of SBE2 enhancer activity leads to reduction in rostral diencephalic *Shh* expression

To examine the role of the SBE2 enhancer during development, this element was deleted using CRISPR/Cas9 (Cong et al., 2013) to remove 1.2 kb (Fig. S1A, B) containing the conserved element. The SBE2 deleted allele, henceforth referred to as *Shh^ΔSBE2^*, was crossed to generate homozygous embryos (*Shh^ΔSBE2/ΔSBE2^*) and also to *Shh* null (*Shh^null^*) mice to generate compound heterozygous embryos (*Shh^ΔSBE2/null^*). Loss of the SBE2 enhancer alone had no effect upon survival as the *Shh*^*ΔSBE2/ΔSBE2*^ mice were viable and fertile and at weaning no reduction in the number of mutant mice was found. Conversely, no live *Shh*^*null/ΔSBE2*^ offspring were recorded by postnatal (P) 14 days (Fig. 1B), these compound heterozygous embryos died perinatally failing to survive beyond P4. Normal ratios were observed at embryonic (E) 17.5 (Fig. 1A).

**Fig. 1.**
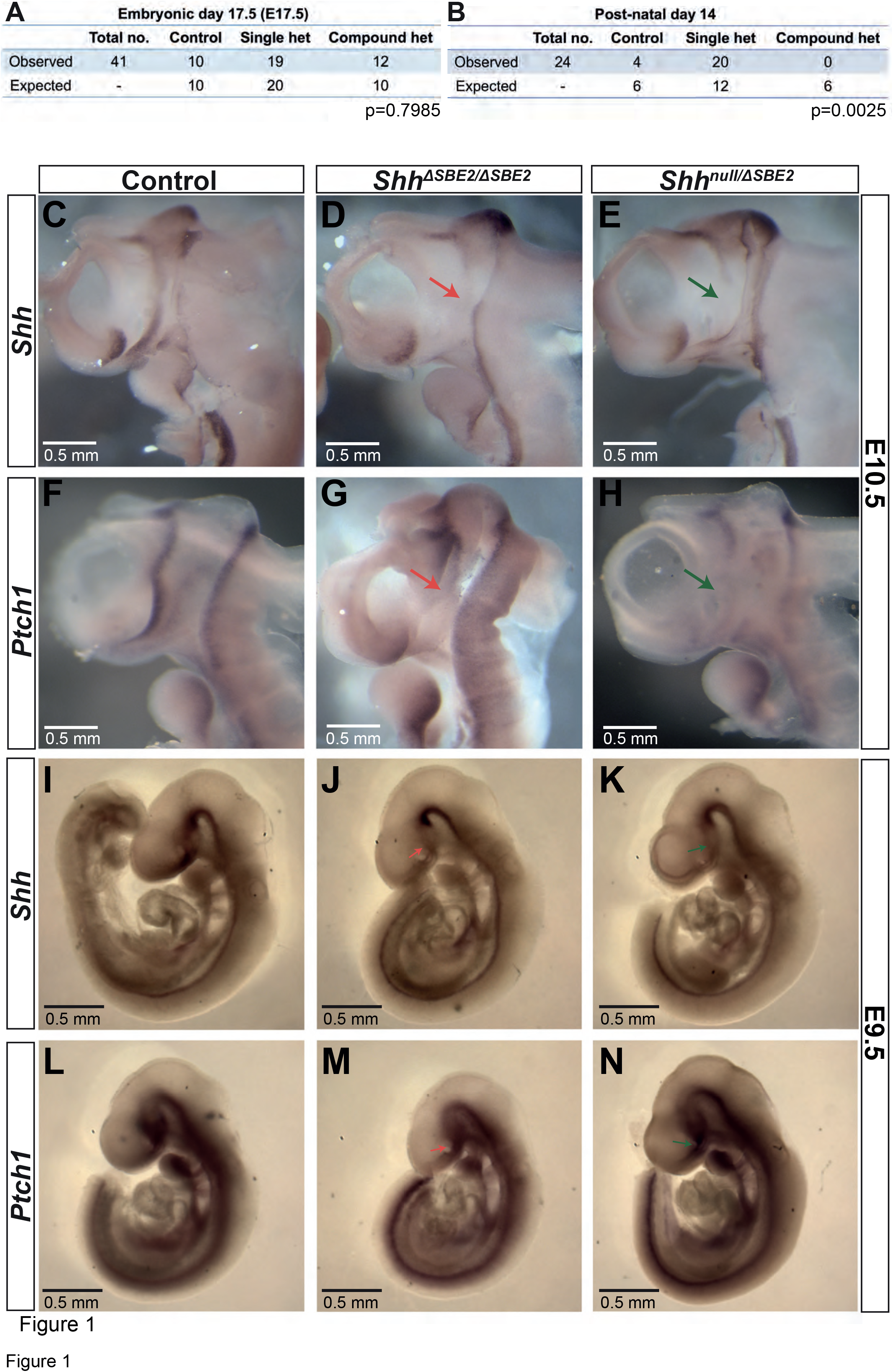
Loss of SBE2 leads to reduced *Shh* activity in the rostral diencephalon. (A-B) Peri and post-natal survival for *Shh*^*null/ΔSBE2*^ compound heterozygotes is shown. Statistical significance was assessed using the Student’s T-test. (C-E) *In situ* hybridisation for *Shh* in E10.5 day heads bisected along the midline is shown for control, *Shh*^*ΔSBE2/ΔSBE2*^ homozygotes and *Shh*^*null/ΔSBE2*^ compound heterozygotes. The red and green arrows point to the rostral diencephalic region where *Shh* expression is absent. (F-H) *In situ* hybridisation for *Ptch1* in E10.5 day bisected heads is shown. Wildtype levels of *Ptch1* expression in rostral diencephalon in control embryos (indicated by black arrow in F), reduced levels in *Shh*^*ΔSBE2/ΔSBE2*^ (indicated by red arrows in G) and undetectable levels in *Shh*^*null/ΔSBE2*^ (indicated by green arrow in H) is shown. Scale bars (C-H) appear in the bottom left hand corner. (I-K) *In situ* hybridisation for *Shh* in E9.5 embryos is depicted for control, *Shh*^*ΔSBE2/ΔSBE2*^ homozygotes and *Shh*^*null/ΔSBE2*^ compound heterozygotes. The red and green arrows point to the rostral diencephalic region where *Shh* expression is lost. (L-N) *In situ* hybridisation for *Ptch1* in E9.5 embryos is shown. The red and green arrows point to the rostral diencephalon. Scale bars (C-N) appear in the bottom left hand corner. (n=3)

The rostral diencephalon, extending from the ZLI to the MGE, is the target for SBE2 expression as predicted by SBE2/reporter gene analysis in transgenic embryos (Jeong *et al*., 2006). *Shh* mRNA expression analysis at E10.5 in *Shh*^*ΔSBE2/ΔSBE2*^ and *Shh*^*null/ΔSBE2*^ embryos revealed undetectable levels of *Shh* expression in the ventral portion of the rostral diencephalon in both crosses (Fig. 1C-E), this region of the diencephalon will give rise to the ventral portion of the tuberal and mammillary hypothalamus (Xie and Dorsky, 2017), and for simplicity will henceforth be referred to as the ventral hypothalamus (VH). The region lacking *Shh* expression in the SBE2 mutant embryos, however, was appreciably more restricted than the region predicted by transgenic analysis, this is likely due to overlapping expression caudally directed by other brain enhancers such as SBE3 and SBE4.

The differences in postnatal viability between *Shh*^*ΔSBE2/ΔSBE2*^ and *Shh*^*null/ΔSBE2*^ suggested that there may be residual, yet undetectable, *Shh* expression in the viable line. To investigate possible low levels of *Shh* expression, patched-1 (*Ptch1*) expression, which is upregulated by hedgehog signalling and is a sensitive readout of *Shh* expression, was examined at E10.5 (Casillas and Roelink, 2018). Similar to the expression pattern observed for *Shh*, *Ptch1* was also downregulated in the VH upon loss of SBE2 activity (Fig. 1F-H). However, in this instance, levels differed between the compound heterozygous and the homozygous embryos. While *Shh*^*null/ΔSBE2*^ embryos showed no detectable levels of *Ptch1* expression in the VH (Fig. 1H, green arrow), the *Shh*^*ΔSBE2/ΔSBE2*^ embryos presented residual levels (Fig. 1G, red arrow). It should be noted that while at E9.5 *Shh* levels are low in the ventral diencephalon in the absence of SBE2 for both mutant conditions (Fig. 1I-K), that *Ptch1* expression pattern in neighbouring tissue is comparable to wildtype (Fig. 1L-N). We suggest that *Ptch1* dependent development is unaffected until after E9.5 and the defects occur after this stage. To determine when VH *Shh* expression is downregulated, a mouse line containing the *LacZ* reporter gene inserted into the *Shh* regulatory domain, SBLac526, was used (Anderson and Hill, 2014) as proxy for *Shh* expression. VH *LacZ* expression was low but detectable up to E13.5 (Fig. S2A-C, red arrow), with expression subsequently absent at E14.5 (Fig. S2D, black arrow). Hence, the consequential role of SBE2 encompasses a developmental window that begins sometime after E9.5 extending to E13.5 when all regulatory activity in the VH is decreasing.

We suspected that presumed residual, low levels of *Shh* expression in the absence of SBE2 were capable of rescuing the postnatal lethality. Accordingly, an additional enhancer was identified in *Shh* expressing neural cells (Harfe et al., 2004) from disected brains. ATAC-seq (Buenrostro et al., 2015) revealed a vertebrate conserved sequence element (position chr5:29230287-29230750 [mm9]), (Fig. S3A) and when used in reporter constructs in transgenic embryos at E10.5 (Fig. S3B-D) showed broad expression activity extending anteriorly from the midbrain into the diencephalon (Fig. S3B-D). Recently, an enhancer whose primary function is to drive expression initially in the prechordal plate, and later throughout the ventral forebrain, called SBE7 (Sagai et al., 2019), was reported at the same chromosomal position. Thus, SBE7 has a primary function in the prechordal plate and secondarily, drives low but sufficient levels throughout the ventral forebrain.

### Loss of SBE2 activity effects midline craniofacial deformities

SBE2 deletion embryos were analysed for craniofacial malformations. Unsurprisingly, given the postnatal viability of *Shh*^*ΔSBE2/ΔSBE2*^ embryos, no gross craniofacial malformations nor deformities in the craniofacial skeletal elements (Fig. S4A-B) were observed. In contrast, at E17.5, *Shh^null/ΔSBE2^* embryos exhibited external head deformities, in which the mutant embryos presented with a more rounded head. To investigate these changes in head shape, control and *Shh*^*null/ΔSBE2*^ mutant embryos were subject to geometric morphometric analysis to deconvolute the skull and test for differences between control and mutant embryos. Centroid size, a proxy of skull scale, revealed that the compound heterozygous mutant embryos tended to have an overall reduction in skull size, though this was not significant (pairwise two-sample Wilcoxon, p=0.49) (Fig. 2A). Accordingly, allometric shape variation due to size, genotype, and litter provided the best regression fit (R^2^=0.83, ANOVA p<0.001). Principal components of the non-allometric shape was also explored. We observed clear, genotype-specific separation of control and mutant mice, with the latter occupying a relatively broader expanse of morphospace (Fig. 2B). In mutants, Principal Component 1 (46.5%) (Fig. 2B) describes a reduction in the anteroposterior axis while dorsoventral and mediolateral axes expanded (Fig. 2C-H). Procrustes variances between control and mutant mice were not significant, however this was presumably due to limitations in statistical power. Indeed, shape differences described by PC1 were significant (Welch Two Sample t-test, p=1.5⨯10^−4^). OPT analysis revealed that, in addition to presenting a perturbed overall head structure, *Shh*^*null/ΔSBE2*^ embryos displayed affected nasal cavities (Fig. 2J,L), which were reduced in length; however, the extent to which the nasal cavities were affected varied between mutant embryos, with the example presented representing a moderate phenotype. Moreover, a region of brain tissue, specifically the midline hypothalamic tissue, was absent in the mutant embryos (Fig. 2I,K).

**Fig. 2.**
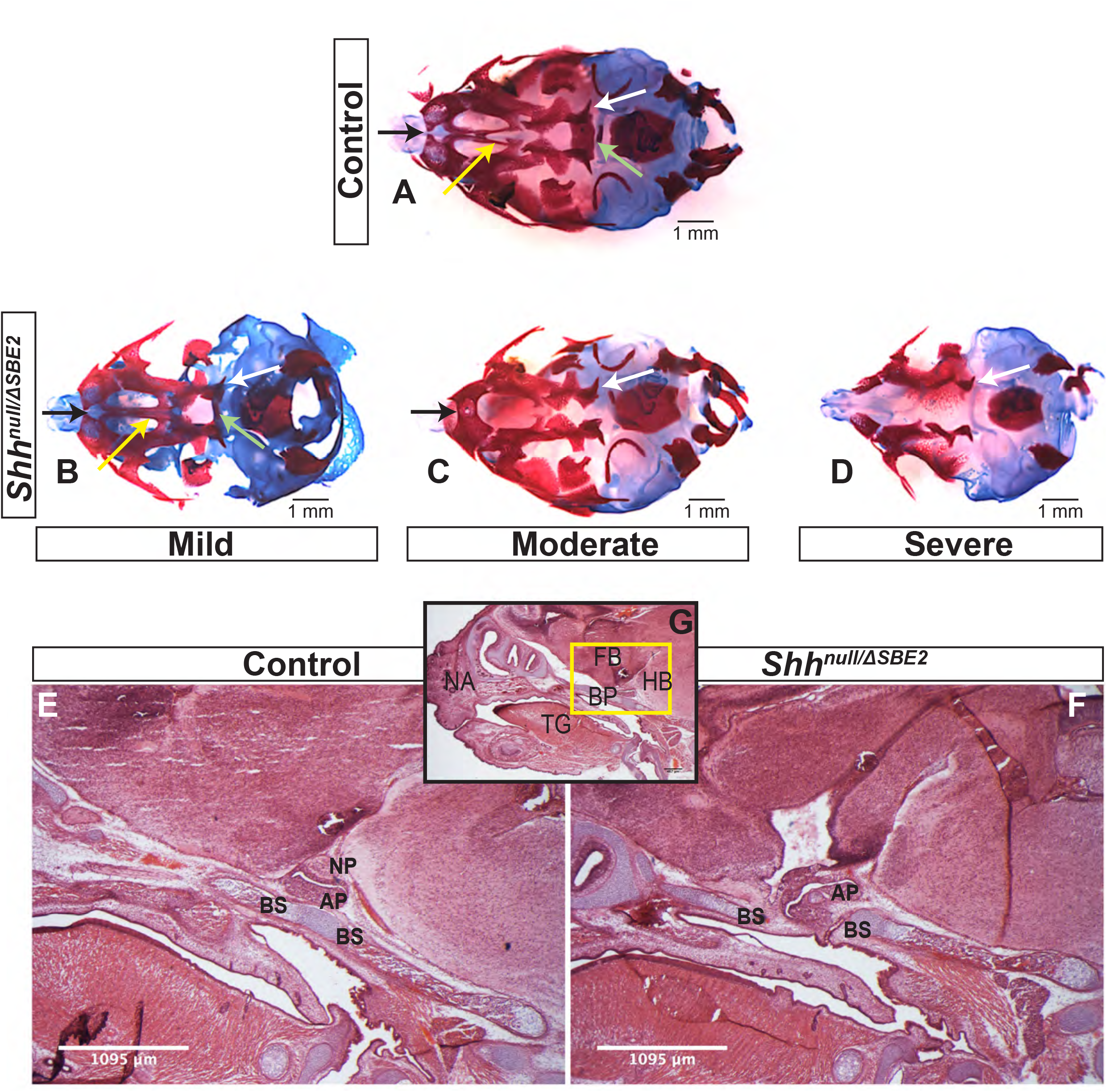
*Shh*^*null/ΔSBE2*^ embryos have an altered head shape and gross deformities. (A) Morphometric analysis of overall head size (log Csize-log centroid size) in control and *Shh*^*null/ΔSBE2*^ heads at E17.5. (B) Morphometric analysis of overall head shape in control and *Shh*^*null/ΔSBE2*^ heads at E17.5. (C-E) Ply surface of a control E17.5 day CT scanned skull, from a dorsal, lateral and ventral angle. (F-H) Morphed ply of a randomly chosen control E17.5.surface, morphed based on the average of *Shh^null/ΔSBE2^* PC1 landmark changes, presented from a dorsal, lateral and ventral angle. (control n=6; mutant n=8). (I-L) OPT scan stills from E17.5 day control and *Shh*^*null/ΔSBE2*^ heads, presenting the midline sagittal plane and the coronal nasal region. Red arrows point to the region of absent hypothalamic tissue seen in *Shh*^*null/ΔSBE2*^ embryos. Red lines serve to demonstrate the reduced nasal cavities found in *Shh^null/ΔSBE2^* embryos. Scale bars are depicted in the bottom left or right hand corner. (n=3)

*Shh*^*null/ΔSBE2*^ embryos displayed midline craniofacial bone deformities with a high degree of variability, which, based on phenotypic severity, were classified into three categories: mild, moderate and severe (Fig. 3A-D). All mutant embryos showed a reduction in the basisphenoid bone, ranging from the presence of only a small remnant (Fig. 3B, pale green arrow) to complete loss (Fig. 3C, D) and in the vomer bone, ranging from an excessive fusion (Fig. 3B, yellow arrow) to complete absence (Fig. 3C, D). In addition, the moderate and severe embryos displayed defects in the pterygoid and maxillary bones (Fig. 3C, D, white arrows). Notably, all embryos showed some midline craniofacial element deformities, which indicates that *Shh* signalling directed by SBE2 is involved in the midline development of the craniofacial bones, and parallels the role *Shh* plays in early midline brain development (Choi et al., 2012).

**Fig. 3.**
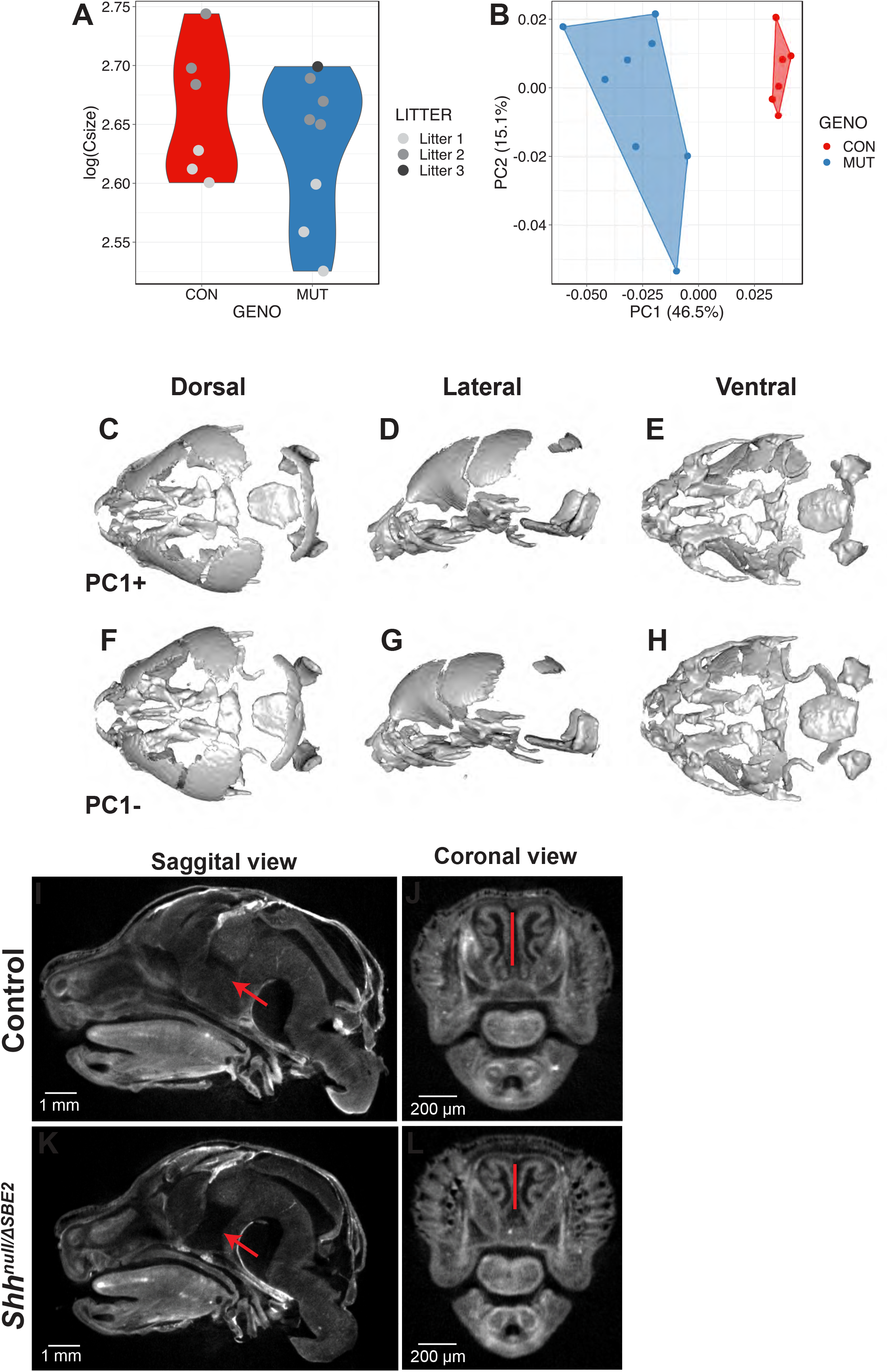
*Shh*^*null/ΔSBE2*^ embryos present deformities in the bones of the cranial vault. (A-D) Dual staining highlighting chondrogenic (blue) and skeletal (red) craniofacial elements in control and *Shh*^*null/ΔSBE2*^ E17.5 heads are shown. Pale green arrows indicate the basisphenoid bone, yellow arrows indicate the vomer bone and the white arrows indicate the pterygoid bone. (E-F) H&E staining of E17.5 day parasagittal cryosections from control and *Shh*^*null/ΔSBE2*^ heads. Black arrows point to the region of the basisphenoid bone which is intercepted by the adenohypophysis in *Shh*^*null/ΔSBE2*^ embryos. A zoomed out view of an E17.5 day head parasagittal cryosection is depicted in (G). The yellow box presents the region of the head depicted in (E-F). AP- adenohypophysis; NP- neurohypophysis; BS- basisphenoid bone; BP- base plate of the skull; NA- nasal prominence; FB- forebrain; HB- hindbrain; TG- tongue. Scale bars are shown in the bottom right or left hand corner. (n=3)

### Mislocalized pituitary and loss of SHH signalling disrupts craniofacial development

Closer examination of the heads of mutant compound heterozygous embryos revealed that the basisphenoid bone, the most posterior defective cranial bone, was intercepted by the adenohypophyseal pituitary lobe (labelled AP in Fig. 3E, F). This lobe of the pituitary was misplaced and formed in a more anterior position, growing within the region where the basisphenoid will form (the basis for the mislocalization is discussed below). Developmental timing of this event suggests that the adenohypophysis disrupts this region before chondrogenesis initiates, which indicates that the adenohypophyseal tissue likely presents a physical barrier to basisphenoid development. The phenotypic variability may correlate with the extent to which the pituitary lobe intercepted the basisphenoid bone in the mutant embryos.

Further investigations into the effect of SBE2 directed expression on craniofacial development focused on mesenchymal *Ptch1* expression in the vicinity of the forebrain. In control embryos at E12.5, *Ptch1* expression was detected within a narrow layer of mesenchyme located directly adjacent to the floor of the diencephalon where *Shh* was expressed (Fig. 4A-B). COL2A1, a type 2 collagen, is expressed during early cartilage formation (Hissnauer et al., 2010) and co-localised with *Ptch1* in control embryos (Fig. 4E-F), implicating that the underlying facial mesenchyme is under the influence of SHH and is committed towards cartilage formation. At the same stage in the *Shh*^*null/ΔSBE2*^ embryos, *Ptch1* expressing cells were absent in the underlying mesenchyme (Fig. 4C-D) and only a thin lining of cells expressing COL2A1 was seen (Fig. 4G-H). The loss of *Ptch1* and COL2A1 staining was found to be independent of the physical disruptive effects caused by mislocalisation of the neurohypophysis, as the absence of both markers was evident in regions of the cranial base which lie anterior to disrupted neurohypophyseal tissue detected in the mutant embryos (Fig. 4C,G). Thus, SBE2 mediated *Shh* expression is instrumental in ensuring adequate cell response and bone formation in the sub– diencephalic mesenchyme along the midline of the palatal floor, and reduction of this cell population is critical for the midline craniofacial phenotype.

**Fig. 4.**
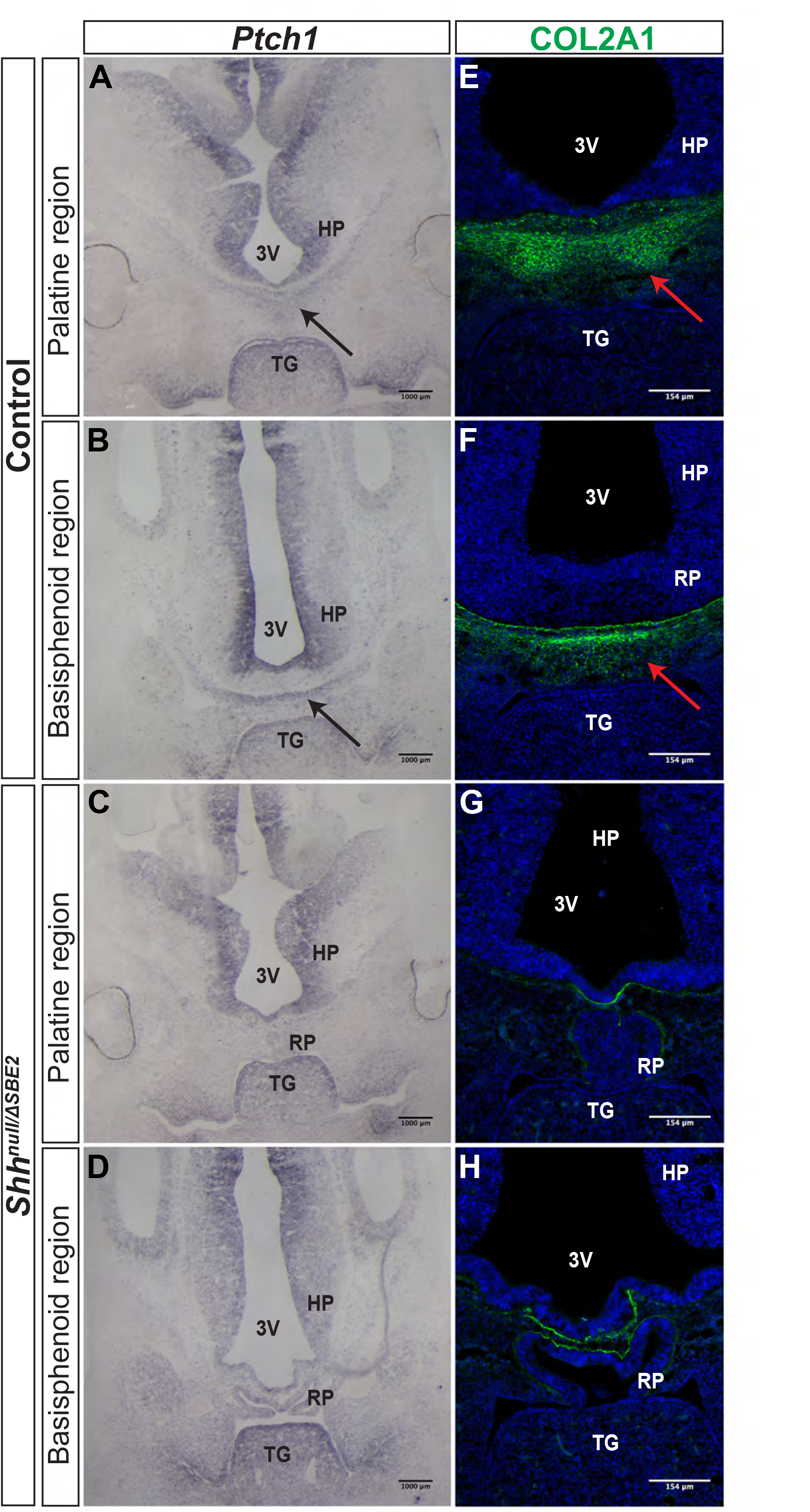
Bone formation in the viscerocranium of *Shh*^*null/ΔSBE2*^ embryos. (A-D) *In situ* hybridisation for *Ptch1* in E12.5 day coronal cryosections of control and *Shh*^*null/ΔSBE2*^ embryos. (A&C) present the palatine region and (B&D) present the basisphenoid region. Black arrows point to the region of *Ptch1* expression seen below the hypothalamic tissue which is absent in *Shh*^*null/ΔSBE2*^ embryos. (E-H) Immunofluorescent staining for COL2A1 in E12.5 day coronal cryosections of control and *Shh^null/ΔSBE2^* embryos. (E&G) present the palatine region and (F&H) present the basisphenoid region. Red arrows point to the region of COL2A1 expression seen below the hypothalamic tissue which mostly is absent in *Shh*^*null/ΔSBE2*^ embryos. 3V- third ventricle; HP- hypothalamus; TG- tongue; RP- Rathke’s pouch. (n=3)

### *Shh*^*null/ΔSBE2*^ embryos display mislocalised and deformed pituitary lobes due to disruption of the AP hypothalamic boundary

Pituitary lobe development was analysed at stages E11.5, E13.5 and E15.5. In *Shh*^*null/ΔSBE2*^ mutant embryos at E11.5, Rathke’s pouch (RP), which will give rise to the adenohypophyseal lobe, and the infundibulum, which will give rise to the neurohypophyseal pituitary lobe, were both ectopically shifted ventrally (Fig. S5 A, B). These findings reflect the previous observations (Zhao et al., 2012) in conditional *Shh* deletion mice, when recombination was driven by the SBE2 enhancer (designated SBE2-Cre Δ*Shh*) an anterior shift in the pituitary lobes was observed. At E13.5, RP was severely malformed in the *Shh*^*null/ΔSBE2*^ mutant embryos, while the infundibulum did not separate from the neuroectodermal tissue to the same degree as control counterparts (Fig. 5A, C) and remained ectopically located at E15.5 (Fig. S5C, D).

**Fig. 5.**
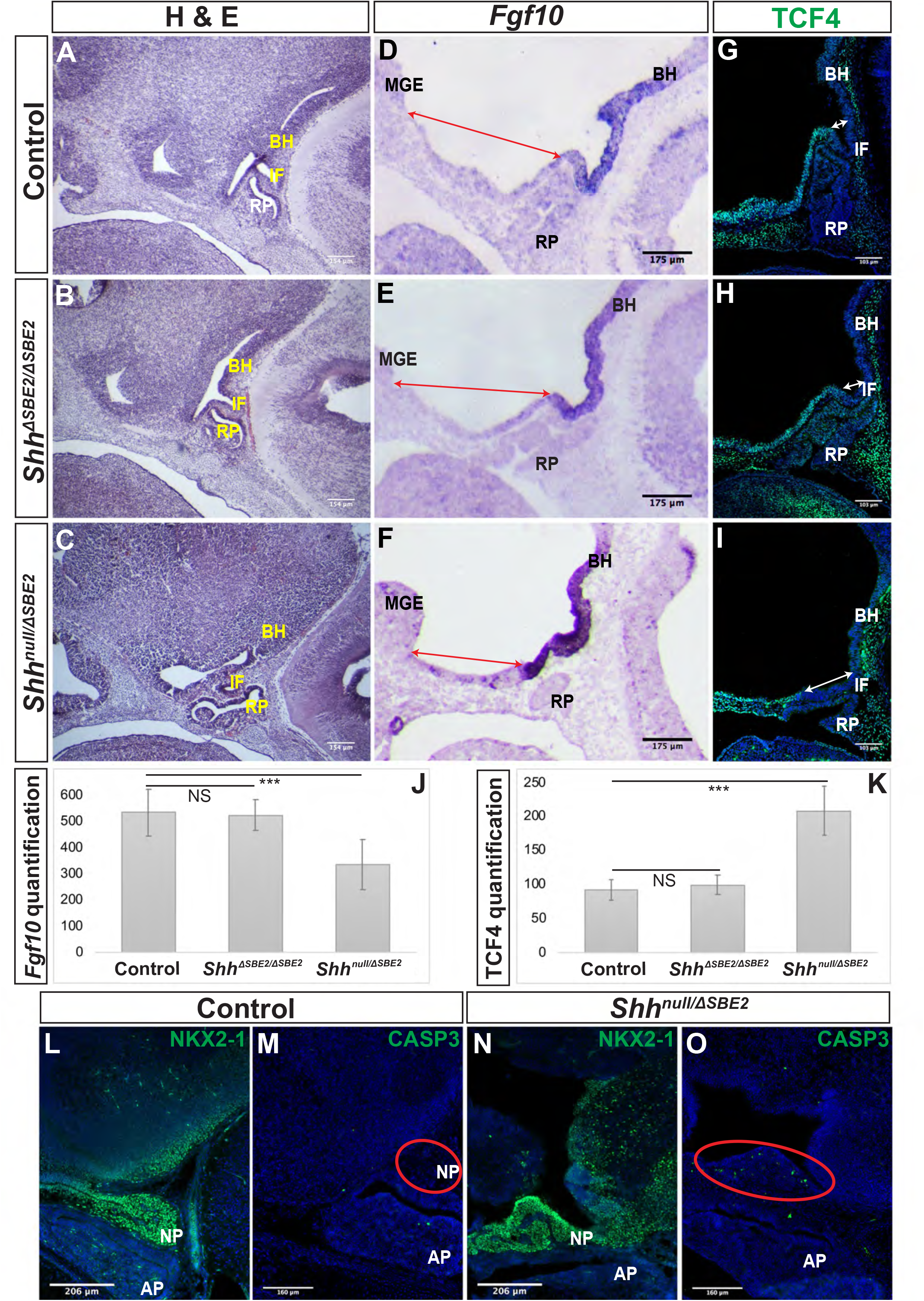
The AP hypothalamic boundary is shifted ventrally in *Shh*^*null/ΔSBE2*^ embryos. (A-C) H&E staining of E13.5 day sagittal cryosections of control, *Shh*^*ΔSBE2/ΔSBE2*^ and *Shh*^*null/ΔSBE2*^ embryos. (D-F) *In situ* hybridisation for *Fgf10* in E11.5 day sagittal cryosections of control, *Shh*^*ΔSBE2/ΔSBE2*^ and *Shh*^*null/ΔSBE2*^ embryos. Red arrows represent the distance between the MGE and the anterior boundary of *Fgf10* expression. (G-I) Immunofluorescent staining for TCF4 in E11.5 day sagittal cryosections of control, *Shh*^*ΔSBE2/ΔSBE2*^ and *Shh*^*null/ΔSBE2*^ embryos. White arrows represent the distance between the posterior infundibulum and the posterior boundary of TCF4 expression. (J)- Quantification of the distance depicted by the red arrows in (D-F) in control, *Shh*^*ΔSBE2/ΔSBE2*^ and *Shh*^*null/ΔSBE2*^ embryos. Statistical significance was assessed using the Student’s T-test. Error bars represent median ± s.e.m. p***<0.001.(K)- Quantification of the distance between depicted by the white arrows in (G-I) in control, *Shh*^*ΔSBE2/ΔSBE2*^ and *Shh*^*null/ΔSBE2*^ embryos. Statistical significance was assessed using the Student’s T-test. Error bars represent median ± s.e.m. p***<0.001. (L & N) Immunofluorescent staining for NKX2-1 in E17.5 day sagittal cryosections of control and *Shh*^*null/ΔSBE2*^ embryos. (M & O) Immunofluorescent staining for CASP3 in E17.5 day sagittal cryosections of control and *Shh*^*null/ΔSBE2*^ embryos. Red circles demarcate the region of cell death apparent in mutant embryos. NS- not significant. BH- basal hypothalamus; IF- infundibulum; RP- Rathke’s pouch; MGE- medial ganglionic eminence. Scale bars are shown in the bottom left hand corner. (n=3)

Interestingly, at E17.5 the neurophypophysis appeared absent in the *Shh*^*null/ΔSBE2*^ mutant embryos (Fig. 3E, F), yet staining for markers of neurohypophyseal tissue revealed that this was due to the lack of separation of the neurohypophysis from the neuroectodermal tissue of origin (Fig. 5L, N). Moreover, caspase-3 (CASP3) staining revealed that this remnant neurohypophyseal tissue appeared to be undergoing apoptosis (Fig. 5M, O) indicating that inadequate neurohypophyseal partitioning likely led to tissue degeneration. Despite the fact that both hypophyseal lobes were misspecified in the *Shh*^*null/ΔSBE2*^ embryos early hypophyseal patterning was unperturbed (Fig. S6C, F, I, L). These findings agree with observations by Zhao et al., (2012) in SBE2-Cre Δ*Shh* mice, where pituitary markers, such as LHX3, were unperturbed despite hypophyseal misplacement. However, whilst SBE2-Cre Δ*Shh* mice displayed multiple invagination sites for RP, only one site was observed in *Shh*^*null/ΔSBE2*^ embryos. To exclude the possibilty that the observed pituitary malformation and mislocalisation, specifically that of the adenohypophysis, were due to a developmental delay, expression of adenohypophyseal markers that display stage-specific restricted expression within the lobe were analysed. ISL1 is expressed throughout the early developing adenohypophysis but later becomes restricted to the ventral most portion of the lobe (Castinetti et al., 2015; Ericson et al., 1998). This same restriction of expression was seen for TBX3, which was expressed broadly throught the developing adenohypophysis, but by E12.5 expression was restricted to the ventral aspect of the lobe (Pontecorvi et al., 2008). For both ISL1 (Fig, S7A-B) and TBX3 (Fig. S7C-D) expression in the adenohypophysis at E12.5 in *Shh*^*null/ΔSBE2*^ embryos was found to reflect the pattern seen in control embryos, with expression only detected in the ventral most portion, indicating that the adenohypophyseal lobe was not developmentally delayed.

While patterning and early neuronal marker expression within the pituitary lobes appeared normal, at later developmental stages, E17.5, neuronal populations within varied nuclei of the VH, which establish connections with the pituitary lobes, were disrupted. Both the tyrosine hydroxylase (TH1) (Fig. S8A-B) and the vasopressin neuronal population (Fig. S8E-F) of the paraventricular nucleus (PVN) were lost in *Shh*^*null/ΔSBE2*^ embryos. Additionally, the oxytocin neuronal population of the PVN was absent while the population found within the arcuate nucleus (ARC) was either misspecified or mislocalised (Fig. S8C-D). To confirm that the loss of PVN and ARC neuronal populations did not result from a developmental delay in hypothalamic neuronal specification, expression of oxytocin and TH1 was analysed at E15.5. At E15.5 oxytocin expression was present in the PVN of the control embryos (Fig. S9A, white arrow) but absent from *Shh*^*null/ΔSBE2*^ embryos (Fig. S9B). Similarly, expression of TH1 was detected in the ARC of control embryos (Fig. S9C, yellow arrow), yet no expression was detected in *Shh*^*null/ΔSBE2*^ embryos (Fig. S9D). This indicates that the lack of specific gene expression in hypothalamic neuronal populations at E15.5 persists for at least two additional days up to E17.5, arguing for a loss of the neuronal population and not a developmental delay in neuronal specification. These results indicate that, in addition to the disruption of hypophyseal structure and location in development, there is also disruption of hypothalamic neuronal populations, demonstrating that SBE2 activity is vital for the role of *Shh* in hypophyseal and hypothalamic neuronal development. Interestingly, the hypothalamic populations found to be affected are known to establish connections with and regulate the hypophysis, indicating that *Shh* plays a role in coordinating hypophyseal development and subsequent neuronal regulation. The disruption of these late neuronal populations in *Shh*^*null/ΔSBE2*^ embryos revealed that early loss of SBE2 directed *Shh* activity in the VH has late effects upon development in the hypothalamus, at timepoints at which *Shh* expression is no longer present in the tissue in question (Fig. S2).

In *Shh*^*ΔSBE2/ΔSBE2*^ mutant embryos, neither the adenohypophysis nor the neurohypophysis were ectopically located and both lobes had separated from the tissue of origin E13.5 (Fig. S4). Additionally, no defects were observed in early marker expression (Fig. S6B, E, H, K); however, mild deformities were detected predominantly in the structure of the lateral portion of adenohypophyseal lobe in *Shh*^*ΔSBE2/ΔSBE2*^ mutant embryos (Fig. S4C-K).

Distinct anterior-posterior expression domains are established along the midline of the hypothalamus by E9.5. The posterior hypothalamic domain expresses factors including *Bmp4* and *Fgf10* whilst the anterior domain expresses *Six6*, *Tcf4* and *Shh*. Mutual inhibition between these two gene expression domains exists, whereby in the absence of one domain the opposing domain will expand to occupy the hypothalamic space (Ericson *et al*., 1998; Takuma *et al*., 1998). The infundibulum expresses the posterior markers *Bmp4* and *Fgf10* and is specified dorsally to the anterior hypothalamic tissue, at the AP transition zone (Ferrand, 1972; Kawamura and Kikuyama, 1995; Takuma *et al*., 1998; Treier *et al*., 1998). Inductive cues from the ventral hypothalamic tissue play a role in orchestrating the formation of RP (Schwind, 1928; Takuma *et al*., 1998). As RP and the infundibulum were both shifted ventrally in the *Shh*^*null/ΔSBE2*^ mutant embryos (Fig. 5C), verification of any other associated defects in the establishment of the AP hypothalamic boundary were sought. The posterior boundary of expression of TCF4 was shifted ventrally in the mutant embryos (Fig. 5I) whereas the anterior boundary of expression of *Fgf10* was shifted ventrally, expanding expression (Fig. 5F). The shift observed for both the anterior and posterior markers was found to be significant in the *Shh*^*null/ΔSBE2*^ mutant embryos (Fig. 5J, K). These results indicated that there is a shift in the AP domain boundary with a reduction in anterior markers and gain of posterior markers. In the *Shh*^*ΔSBE2/ΔSBE2*^ mutant embryos no shift in expression of TCF4 (Fig. 5H) or *Fgf10* (Fig. 5E) was observed, correlating with correct positioning of the pituitary lobes (Fig. 5C). These results suggest that shift of the pituitary lobes occurs as a result of the aberrant specification of the AP boundary, which does not appear to be sensitive to reduced levels of SHH shown in *Shh*^*ΔSBE2/ΔSBE2*^ embryos (Fig. S4C-F).

### Loss of SBE2 enhancer activity leads to defects in hypothalamic patterning in a concentration dependent manner

Much of our knowledge about DV hypothalamic patterning is informed by what is known regarding DV neural tube patterning. In the developing neural tube, neural progenitors acquire distinct transcriptional identities based on length of exposure to patterning signals and DV positioning (Balaskas et al., 2012; Dessaud et al., 2007, 2010). Indeed it is the opposing action of the dorsal neural tube signals *Wnts* and BMPs and the ventral neural tube signal *Shh* that drive patterning along this axis (Burbridge et al., 2016). Subsequently, expression of distinct combinations of transcription factors drives the differentiation of the neural progenitors towards specific neural fates (Dessaud et al., 2008). In the developing hypothalamus *Shh* is initially expressed from the prechordal mesoderm; it is subsequently expressed along the floor plate and plays a role in inducing ventral hypothalamic fate (Burbridge et al., 2016; Corman et al., 2018). In contrast, BMPs are expressed from the roof of the telencephalon, regulating alar hypothalamic development (Burbridge et al., 2016). Given that *Shh* signalling from the hypothalamic floor plate was reduced in the *Shh*^*ΔSBE2/ΔSBE2*^ and *Shh*^*null/ΔSBE2*^ mutant embryos, DV hypothalamic patterning was examined. Genes whose expression are responsive to SHH levels in the neural tube were of particular interest. No differences were detected between wild-type, *Shh*^*ΔSBE2*/+^ and *Shh*^*null*/+^ embryos (Fig. S10), indicating that a 50% reduction in *Shh* levels has no effect upon DV patterning marker expression.

Expression of PAX6, known to the expressed within the alar hypothalamus, and the opposing basal hypothalamic marker NKX2*-2* (Croizier et al., 2015; Guillemot, 2007) were examined. NKX2-2 expression commences early in hypothalamic development, with expression detected in the basal hypothalamic region which borders the alar zone. Later in development, this expression pattern alters within the VH, whereby expression has forked into two bilateral stripes; one that occupies the transition zone of the alar-basal hypothalamus extending into the alar hypothalamus itself, and a second stripe that resides within the basal hypothalamic territory of the VH (Burbridge et al., 2016). At E13.5, this bilateral stripe expression pattern was observed in control embryos (Fig. 6A). However, in *Shh*^*ΔSBE2/ΔSBE2*^ mutants the dorsal VH stripe, within the basal hypothalamus, presented patchy expression (Fig. 6B). The *Shh*^*null/ΔSBE2*^ mutant embryos were more severely affected, with both the alar and basal VH stripes absent (Fig. 6C). This data indicate that low levels of *Shh* expression can induce some NKX2-2 expression in *Shh^ΔSBE2/ΔSBE2^*, however, these low levels are not sufficient to either specify or maintain the entirety of the NKX2-2 population.

**Fig. 6.**
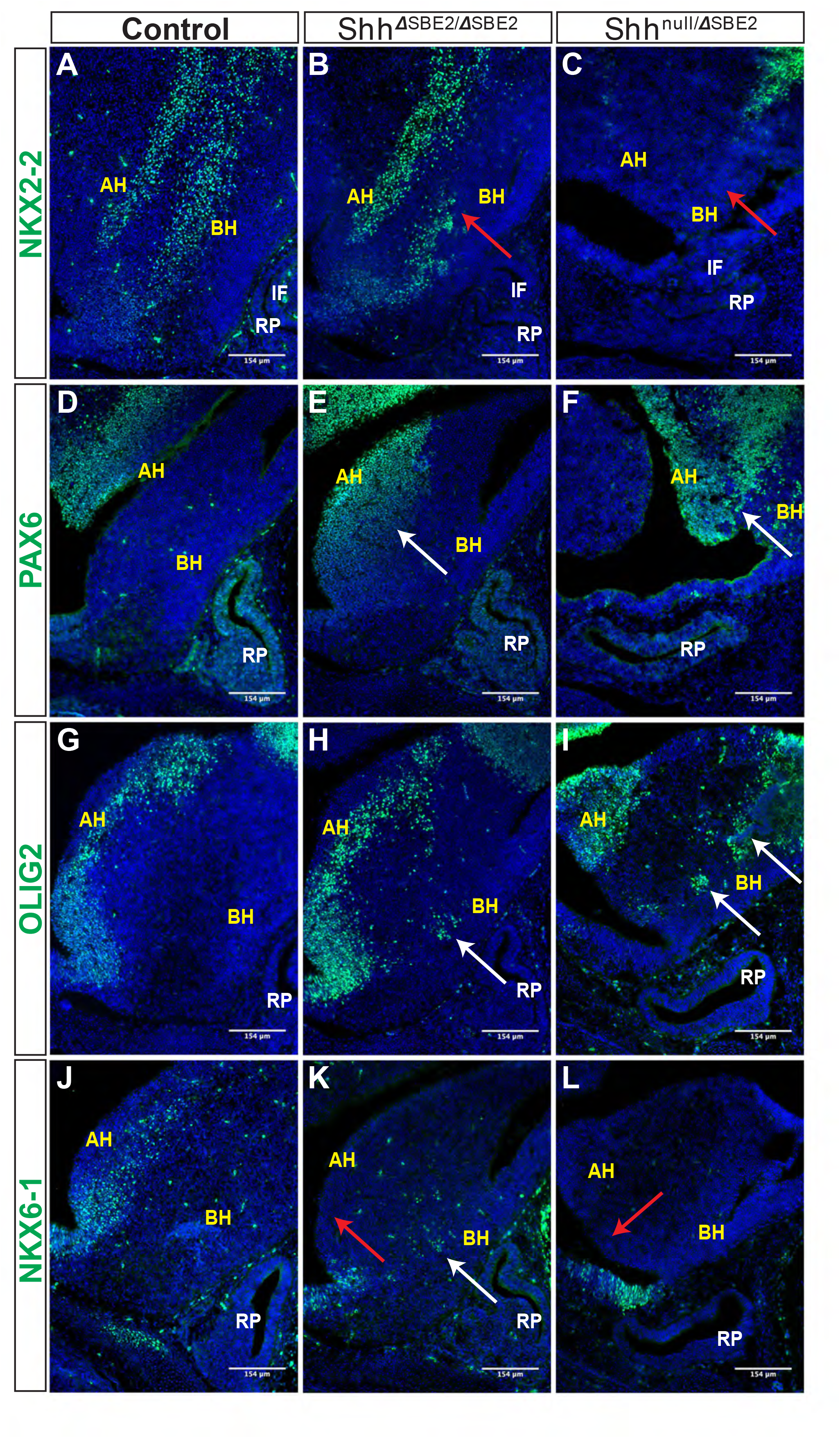
DV hypothalamic patterning is affected by loss of SBE2. (A-C) Immunofluorescent staining for NKX2-2 in E13.5 day ventral hypothalamic (VH) parasagittal cryosections of control, *Shh*^*ΔSBE2/ΔSBE2*^ and *Shh*^*null/ΔSBE2*^ embryos. Red arrows point to the region of NKX2-2 expression in the basal hypothalamus which is reduced or absent upon removal of SBE2 enhancer function. (D-F) Immunofluorescent staining for PAX6 in E13.5 day VH parasagittal cryosections of control, *Shh*^*ΔSBE2/ΔSBE2*^ and *Shh*^*null/ΔSBE2*^ embryos. White arrows point to the region of ectopic PAX6 expression in the BH which is seen upon removal of SBE2 function. (G-I) Immunofluorescent staining for OLIG2 in E13.5 day VH parasagittal cryosections of control, *Shh*^*ΔSBE2/ΔSBE2*^ and *Shh*^*null/ΔSBE2*^ embryos. White arrows point to the ectopic regions of expression of OLIG2 in the BH which are seen upon removal of SBE2 function. (J-L) Immunofluorescent staining for NKX6-1 in E13.5 day VH parasagittal cryosections of control, *Shh*^*ΔSBE2/ΔSBE2*^ and *Shh*^*null/ΔSBE2*^ embryos. Red arrows point to the region of NKX6-1 expression in the AH which is absent upon removal of SBE2 function whilst white arrows point to the region of ectopic expression in the BH seen in *Shh*^*ΔSBE2/ΔSBE2*^ embryos. AH- alar hypothalamus; BH- basal hypothalamus; IF- infundibulum; RP- Rathke’s pouch. Scale bars are shown in the bottom right hand corner. (n=3)

PAX6 is a neural tube and hypothalamic marker, whose expression pattern is known to be restricted to the alar hypothalamus (Burbridge et al., 2016) and, as expected, E13.5 control embryos displayed PAX6 expression solely within the alar portion of the VH (Fig. 6D). In *Shh*^*ΔSBE2/ΔSBE2*^ mutant embryos there was a dorsal expansion of PAX6 expression into the basal hypothalamic region in the embryos (Fig. 6E). In *Shh*^*null/ΔSBE2*^ embryos, this dorsal expansion of PAX6 expression was more severe, with large patches of PAX6 expression detected in the basal hypothalamus of the mutant embryos (Fig. 6F). These results indicate that *Shh* signalling from the basal hypothalamus is required to restrict PAX6 expression solely to the alar region, and that low levels of *Shh* activity, those found in basal hypothalamus of *Shh*^*ΔSBE2/ΔSBE2*^ mutants, are not sufficient to exclude PAX6 activity from this region.

These results for NKX2-2 and PAX6 within the VH, upon loss of SHH signalling activity, mirror what was found by Corman and colleagues (2018). However, these data demonstrate a novel dose dependent effect of SHH signalling whereby these transcription factors are not only able to respond to the presence, or indeed absence, of *Shh* signalling, but are also sensitive to the amount of *Shh*, with the levels of *Shh* activity being key for theestablishment of hypothalamic signaling domains.

Expression of the transcription factor OLIG2 is induced by SHH (Dessaud et al., 2008), and in control embryos was detected in the alar hypothalamus in a narrow region along the edge of the third ventricle (Fig. 6G). In *Shh*^*ΔSBE2/ΔSBE2*^ mutant embryos, however, the OLIG2 positive cells showed a subtle dorsal shift away from the edge of the third ventricle and additional expressing cells were detected in the basal hypothalamus (Fig. 6H). In *Shh*^*null/ΔSBE2*^ mutant embryos, OLIG2 positive staining cells had dramatically shifted into the basal portion of the hypothalamus, with large areas of positive staining cells detected (Fig. 6I). These data indicated that *Shh* activity acts to ensure adequate OLIG2 expression within the alar hypothalamus, whereupon reduction of SHH allows for OLIG2 expression within the basal hypothalamus. Unexpectedly, OLIG2 was found to be expanded into ectopic regions of the VH in mutant embryos rather than gradually reduced with increasing loss of SHH activity. However, Balaskas et al., 2012, demonstrated in the neural tube that, whilst NKX2-2 and OLIG2 are both equally responsive to SHH activity, higher levels of signalling are required for NKX2-2 expression as it is repressed by both PAX6 and OLIG2, creating a gene regulatory network between these three transcription factors. Thus, in this instance, the residual levels of *Shh* expression directed by SBE7 in *Shh*^*null/ΔSBE2*^ mutant embryos are likely sufficient to induce OLIG2 expression, yet not NKX2-2 due to the repressive activity of OLIG2 and PAX6.

NKX6-1, another transcription factor induced by SHH in the neural tube (Dessaud et al., 2008), was expressed in the alar hypothalamus in a narrow region along the edge of the third ventricle in sagittal and parasagittal regions (Fig. 6J); but in contrast to OLIG2, these cells were only detected along the basal lateral edge of the third ventricle. In *Shh*^*ΔSBE2/ΔSBE2*^ mutant embryos, NKX6-1 expression along the basal lateral edge of the third ventricle was undetectable (Fig. 6K), however, isolated groups of expressing cells were detected in the basal hypothalamus (Fig. 6K). *Shh*^*null/ΔSBE2*^ mutants showed complete loss of the region of staining along the basal lateral edge with no additional regions of expression observed (Fig. 6L). NKX6-1 appears to require low levels of *Shh* expression to be induced, levels which are normally found along the edge of the third ventricle. In *Shh*^*ΔSBE2/ΔSBE2*^ mutants the levels of *Shh* expression are reduced within the basal hypothalamus to a level that allows for ectopic expression of NKX6-1 within this region. In *Shh*^*null/ΔSBE2*^ mutants the complete loss of the expressing cells from the edge of the third ventricle indicate that the levels of *Shh* signalling activity within the VH, directed by SBE7, are insufficient to induce expression of NKX6-1. These data demonstrate a novel, previously undescribed role for SHH activity in the regulation of NKX6-1 expression within the hypothalamus, which mirrors the role described in the neural tube (Sander et al., 2000). Thus, providing further evidence that the signalling network which operates in the neural tube has been co-opted in hypothalamic development.

For all hypothalamic markers analysed, the extent of changes in expression correlate to the scale of alteration in levels of *Shh* expression, suggesting that *Shh* signalling is functioning in a concentration dependent manner to induce ventral patterning in the hypothalamus. The data described not only present novel examples of patterning genes regulated by SHH activity, as is the case for NKX6-1, but also expand on data presented by Corman *et al.* (2018). We further demonstrate that, not only do these genes require SHH activity to be adequately expressed, but they are capable of responding to varying levels of SHH. This concentration dependent effect mirrors that seen for *Shh* in the neural tube, and involves many of the same key players, indicating that *Shh* is acting as a morphogen within the VH to regulate tissue establishment within this critical brain region.

### SBE2 enhancer activity controls downstream effects upon hypothalamic neuronal populations

We next sought to assess the effects of disrupted DV patterning on neuronal populations. We first verified that there were no detectable differences in marker expression between wild-type and heterozygotes, *Shh*^*ΔSBE2*/+^ and *Shh*^*null*/+^ embryos (Fig. S11), demonstrating that a 50% reduction in *Shh* levels has no gross effect upon DV neuronal population specification.

To examine the effect of disrupted DV patterning upon the establishment of the different ventral hypothalamic neuronal populations, we first analysed expression of the broad ventral hypothalamic neuronal marker NKX2-1 (Ohyama et al., 2005). In control embryos at E13.5, NKX2-1 positive cells were detected across the entirety of the basal hypothalamic domain (Fig. 7A). In *Shh*^*ΔSBE2/ΔSBE2*^ mutant embryos, the region of NKX2-1 expressing cells was reduced and embedded within smaller domains intermingled with patches devoid of staining (Fig. 7B). In the *Shh*^*null/ΔSBE2*^ mutant the region of expression was further reduced with larger and more numerous patches of negative cells observed (Fig. 7C). These results indicate that whilst SHH activity is required for proper induction of the NKX2-1 neuronal population another factor must also sustain the remnant expression seen in mutant embryos.

**Fig. 7.**
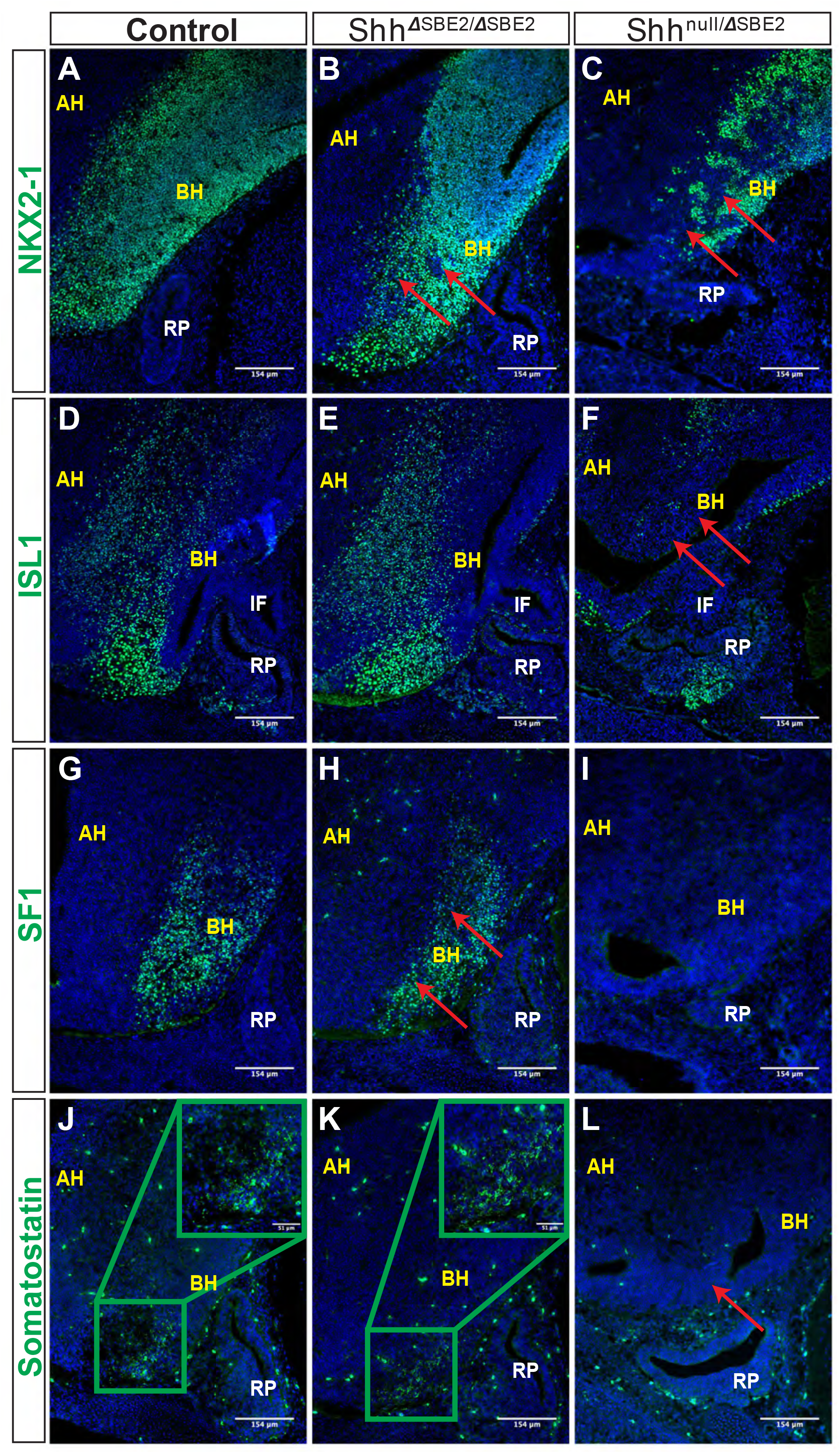
DV hypothalamic neuronal populations are affected by loss of SBE2. (A-C) Immunofluorescent staining for NKX2-1 in E13.5 day VH parasagittal cryosections of control, *Shh*^*ΔSBE2/ΔSBE2*^ and *Shh*^*null/ΔSBE2*^ embryos. Red arrows point to the regions devoid of NKX2-1 expression which are found upon removal of SBE2 function. (D-F) Immunofluorescent staining for ISL1 in E13.5 day VH parasagittal cryosections of control, *Shh*^*ΔSBE2/ΔSBE2*^ and *Shh*^*null/ΔSBE2*^ embryos. Red arrows point to the regions of the BH where ISL1 is mostly absent in *Shh*^*null/ΔSBE2*^ embryos. (G-I) Immunofluorescent staining for SF1in E13.5 day VH parasagittal cryosections of control, *Shh*^*ΔSBE2/ΔSBE2*^ and *Shh*^*null/ΔSBE2*^ embryos. Red arrows point to the region of the BH where SF1 expression is reduced or absent upon removal of SBE2 function. (J-L) Immunofluorescent staining for somatostatin in E13.5 day VH parasagittal cryosections of control, *Shh*^*ΔSBE2/ΔSBE2*^ and *Shh*^*null/ΔSBE2*^ embryos. Red arrows point to the region of the BH where somatostatin is lost in *Shh*^*null/ΔSBE2*^ embryos. A zoomed in view of the region of somatostatin expression is presented in the green boxes in the upper right hand corner of (J & K). AH- alar hypothalamus; BH- basal hypothalamus; IF- infundibulum; RP- Rathke’s pouch. Scale bars are shown in the bottom right hand corner. (n=3)

ISL1 is a melanocortinogenic neuronal marker, for which positive cells are detected throughout the hypothalamus (Kim et al., 2015). In control embryos positive staining was detected throughout the basal hypothalamic region, as expected (Fig. 7D). No differences in staining were detected between control and *Shh*^*ΔSBE2/ΔSBE2*^ mutant embryos (Fig. 7E), however, in *Shh*^*null/ΔSBE2*^ mutants there was substantial loss of positive staining cells in the basal hypothalamus (Fig. 7F).

We additionally verified whether neurons that are restricted to specific nuclei of the basal hypothalamus were affected upon removal of SBE2 activity. Steroidogenic factor 1 (SF1) positive neurons are localised to the ventromedial hypothalamus (VMH) and are involved in the regulation of glucose metabolism and energy balance (Choi et al., 2013). In control embryos, as expected, SF1 positive neurons were detected in the developing VMH of the basal hypothalamus (Fig. 7G). In *Shh*^*ΔSBE2/ΔSBE2*^ mutant embryos the region of the basal hypothalamus expressing SF1 was reduced (Fig. 7H), while in *Shh*^*null/ΔSBE2*^ mutant embryos there was a complete loss of SF1 positive neurons (Fig. 7I). Somatostatin (SOM) expressing neurons are a subset of GABAergic inhibitory neurons that are detected in varied nuclei of the hypothalamus, including the VMH and the arcuate nucleus, which serve to regulate cell proliferation ultimately affecting growth (Urban-Ciecko and Barth, 2016). In control embryos, SOM neurons were detected in the basal hypothalamic region corresponding to the arcuate nucleus at E13.5 (Fig. 7J). No differences were detected in *Shh*^*ΔSBE2/ΔSBE2*^ mutant embryos where comparable staining was detected (Fig. 7K); however, in *Shh*^*null/ΔSBE2*^ mutant embryos no positive staining for SOM was detected in the VH (Fig. 7L). These results indicated that low levels of *Shh* in the VH were sufficient to retain specification of SOM neurons, but higher levels of *Shh* activity were required to sustain and/or specify the SF1 neuronal population in the basal hypothalamus. In the *Shh*^*null/ΔSBE2*^ mutant embryos both of these populations were absent which indicated that early *Shh* activity was essential for the adequate establishment of both the SF1 and SOM neuronal populations of the basal hypothalamus.

Corman *et al.* (2018), have previously demonstrated that loss of *Shh* signalling directed by SBE2 leads to a loss or reduction of neuronal markers. Our data expand on these findings by demonstrating a previously unknown concentration dependent requirement for *Shh* in VH neuronal development whereby some neuronal populations, such as ISL1 and SOM, are only affected when the levels of *Shh* are vastly reduced, as in *Shh*^*null/ΔSBE2*^ mutants. In contrast, the NKX2-1 and SF1 populations are affected by moderate reductions in the levels of *Shh* and further affected by greater loss of *Shh* signalling.

## Discussion

### Secondary enhancer activity rescues SBE2 inactivation

Loss of SBE2 in *Shh*^*ΔSBE2/ΔSBE2*^ homozygous mice did not cause lethality or overt phenotypic effects in the brain and craniofacial regions due to additional enhancer activity most likely from the broadly acting element SBE7. A multistep series of events generate the normal *Shh* expression levels in the ventral diencephalon of which SBE2 activity plays a later role. SBE7 is responsible for early *Shh* expression throughout the prechordal plate a tissue that underlies the developing brain, from as early as E7.5 (Sagai et al., 2019). SBE7, subsequently, drives *Shh* expression at low levels in the ventral midline of the developing forebrain throughout later developmental stages. By E9.5 SBE2 activation, which is dependent on the earlier SBE7 activity (Sagai et al., 2019), contributes to a substantially increased level of expression specifically in the ventral diencephalon.

SBE2 activity does not appear to be obligatory since in its absence the animals are viable and fertile; raising questions about the value of this enhancer to the embryo. In contrast, the deep conservation of this element strongly argues that SBE2 provides fitness to individuals and is crucial across the vertebrate classes. Consequently, we argue that this is not an example of enhancer redundancy that commonly occurs in the mammalian genome (Osterwalder et al., 2018). Our analysis shows that expression of DV patterning genes is sensitive to level of SHH which is perturbed in the absence of SBE2. SBE2 and SBE7 are components of a systematic temporal response required to reach sufficient SHH levels to attain the precise pattern of gene expression. Since further reduction of *Shh* levels (as in the *Shh*^*null/ΔSBE2*^ mutant) causes greater disruption to the expression of hypothalamic marker genes and perinatal lethality, it is reasonable to argue that, in the absence of SBE2, these animals are not fit in the wild. Disruption of neuronal populations seen in *Shh*^*ΔSBE2/ΔSBE2*^ homozygous embryos, such as the reduction in the NKX2-1 and SF1 positive population, may lead to phenotypic effects on physiology, such as reproduction and energy balance. Indeed, specific depletion of the SF1 neuronal population of the VMH is known to be viable, however the metabolic response of skeletal muscle are attenuated, revealing an effect upon metabolism (Fujikawa et al., 2016).

### SBE2 mediated expression links brain and craniofacial development

Heterozygous inactivating mutations in the *SHH* gene in patients cause HPE (Chiang et al., 1996; Ming and Muenke, 2002) while in mice complete loss of *Shh* leads to multiple phenotypic consequences including the absence of brain and craniofacial structures (Chiang et al., 1996). While these observations highlight the links between brain and craniofacial development, the question as to whether *Shh* signalling from the neuroectoderm is directly required for the development of the craniofacial elements has remained largely unanswered. Physical interactions between the brain and face begin early at the initial stages of facial formation (Marcucio et al., 2011). There are two fundamentally different mechanisms by which development of the face and brain may be interrelated; firstly, the developing brain serves as a dynamic architectural foundation upon which the face responds and develops and secondly, signalling directly from the brain to the cranium and face regulates morphogenesis.

In fish, *Shh* signalling from both the oral ectoderm and neural tissue is required for adequate progression of craniofacial development (Wada et al., 2005). In mammalian development, *Shh* signalling from the oral ectoderm, which underlies the cranial base and also neighbours the facial skeletal elements, has been demonstrated to be key in adequate craniofacial development. Depletion of signalling from the oral ectoderm leads to defects in the maxillary and mandibular bones, the palatine bones and the basisphenoid bone among others (Jeong et al., 2004). The present study addressed the role of SBE2 forebrain expression activity on the development of adjacent tissue which would be predicted to form the midline of the cranial floor. Loss of SBE2 in the presence of the *Shh* null allele leads to both lethality and craniofacial deformities. Defects were observed in multiple skeletal elements of the cranial base including the vomer bone and the basisphenoid bone, which resides below the pituitary glands and the hypothalamic tissue, demonstrating that *Shh* signalling from the rostral diencephalon is directly required for adequate craniofacial develop.

### SBE2 affects both AP and DV patterning in the hypothalamus

In addition to the craniofacial defects observed in the SBE2 mutant mouse lines, defects in hypothalamic development were also observed. This is perhaps unsurprising given that the region of *Shh* expression directed by SBE2 is the rostral diencephalon, the tissue from which the hypothalamus originates. However, the mode in which *Shh* regulates hypothalamic patterning and establishment appears to differ depending on the axis considered.

The AP axis was less sensitive to *Shh* levels of expression, whereby low, residual levels of signalling activity in the *Shh*^*ΔSBE2/ΔSBE2*^ mutants were sufficient to sustain normal patterning. In contrast, further reduction of *Shh* expression in the *Shh*^*null/ΔSBE2*^ mutant embryos caused a loss of the anterior hypothalamic identity, which is promoted by *Shh*, in favour of the posterior hypothalamic identity. A shift in the anterior marker TCF4 was observed in the *Shh*^*null/ΔSBE2*^ mutant embryos, which contrasts previous observations (Zhao *et al*., 2012), where a shift in all other AP hypothalamic markers with the exception of TCF4 was observed. This difference likely resides in the different mouse lines used, whereby the use of SBE2-Cre to conditionally knock-down *Shh* activity may lead, firstly, to a temporal delay in the removal of *Shh* activity and, secondly, to a broader removal of *Shh* activity than that solely directed by SBE2. Notably, *Shh* is also required for adequate development of neuronal populations which connect to the hypophyseal lobes and in turn, regulate hormonal release. These results demonstrate a dual role for *Shh* in ensuring hypophyseal function, directly in hypophyseal development and secondarily in regulating innervation.

In contrast to AP patterning, the DV hypothalamic axis was affected by *Shh* activity in a concentration dependent manner. This is apparent in the *Shh*^*ΔSBE2/ΔSBE2*^ mutant embryos where defects in DV patterning, and also subsequent neuronal induction, were observed. These defects were far less severe than those observed in the *Shh*^*null/ΔSBE2*^ mutant embryos, where the disruption to DV development resulted in a complete loss of hypothalamic neuronal populations. The results presented regarding DV hypothalamic patterning are comparable to those described by Corman *et al*. (2018) where the same key neural tube patterning genes and subsequence neuronal regulators are involved. However, we have demonstrated a novel concentration dependent patterning activity of *Shh* in the hypothalamus, which has previously only been described in the neural tube. Moreover, the results demonstrate loss of neuronal populations beyond those of the dorsomedial, ventromedial and arcuate nuclei, additionally implicating neurons of the paraventricular nucleus, which play a key role in hypophyseal regulation. Corman *et al*. (2018) also propose that NKX2-2 is expressed in a domain between dorsal PAX6 and ventral NKX2-1; however, our data suggest that NKX2-1 is expressed broadly across the entire ventral hypothalamic region, with NKX2-2 expressed in a subdomain of the ventral hypothalamus, in regions overlapping with NKX2-1. It is possible that the ventral hypothalamic markers induced by *Shh*, including NKX2-2 are responsible for inducing expression of NKX2-1, whereby ventral hypothalamic markers that are not completely absent in the *Shh*^*null/ΔSBE2*^ mutant embryos, sustain remaining NKX2-1 expression.

Similar to the patterning process in the neural tube, the response of different DV patterning transcription factors to *Shh* was concentration dependent. For example, NKX6-1 was ventrally displaced in the *Shh*^*ΔSBE2/ΔSBE2*^ mutants, much like OLIG2. However, it was lost altogether in *Shh*^*null/ΔSBE2*^ mutant embryos, unlike OLIG2 which was further displaced; indicating that NKX6-1 is more sensitive to loss of *Shh* signalling than OLIG2. Similarly, in the neural tube OLIG2 responds to a wider range of *Shh* signalling levels than NKX6-1. This same variable sensitivity to *Shh* signalling was seen for the neuronal populations, whereby only some of the neuronal population that were disrupted in the *Shh*^*null/ΔSBE2*^ mutant embryos were also affected in the *Shh*^*ΔSBE2/ΔSBE2*^ mutants; for example, NKX2-1 expression required higher levels of *Shh* expression than those needed for ISL1. Moreover, *Shh* was found to only be essential to maintain the SF1 and somatostatin neuronal populations, which were completely absent in *Shh*^*null/ΔSBE2*^ mutants. Thus, early SBE2 directed *Shh* signalling directs subsequent DV patterning in a concentration dependent manner acting as a morphogen within ventral hypothalamic development, in which transcription factor expression patterns are controlled by *Shh* levels which in turn instruct ventral hypothalamic neuronal populations.

### SBE2 coordinates craniofacial development and patterning of the hypothalamus

SBE2 regulates the expression of *Shh* in a region of the diencephalon in which SHH signalling coordinates development of both the diencephalon and the neighbouring tissue responsible for the cranial and facial bones, skeletal elements which are in turn responsible for protecting the neural structures from external exposure. Homologues of the craniofacial bones disrupted in mouse are found in species of all vertebrate classes from fish to mammals (Maddin et al., 2016). In addition, the hypothalamus is an ancient brain centre critical for the production and release of key hormones (Xie and Dorsky, 2017). We suggest that deep in vertebrate evolution SBE2 activity arose as a means to govern VH development and as a consequence of hypothalamus proximity to craniofacial mesenchyme enabled *Shh* signalling to control formation of the bone to ensure its own protection. We have demonstrated that disruptions to coordinated development of the VH and cranial vault in mice leads to phenotypic defects which reflect those seen for HPE patients. These results reveal not only that the interactions between the brain and the face display deeply conserved evolutionary roots, but also show the key importance of mouse models for addressing human disease phenotypes.

## Author Contributions

Conceptualization, REH.; Methodology, LAL, NK and ZCS.; Validation, ZCS and DMK.; Investigation, ZCS, JS, KAG, PSD, LR, MD, JIH, LAL Writing – Original Draft, ZCS and REH; Writing – Review & Editing, DMK., JS and LAL; Visualization, ZCS, KAG, JS and LAL.; Supervision, DMK, JS and REH; Funding Acquisition, DMK and REH

## Acknowledgements

We would like to thank the MRC Central Services for providing DNA sequence support. We would also like to thank the staff at the Evans Building for expert technical assistance. Collaborative work was supported by the MRC and the University of Calgary

