## Supplemental Figures for "A Highly Conserved *Shh* Enhancer Coordinates Hypothalamic and Craniofacial Development"

### *Supplementary Material*

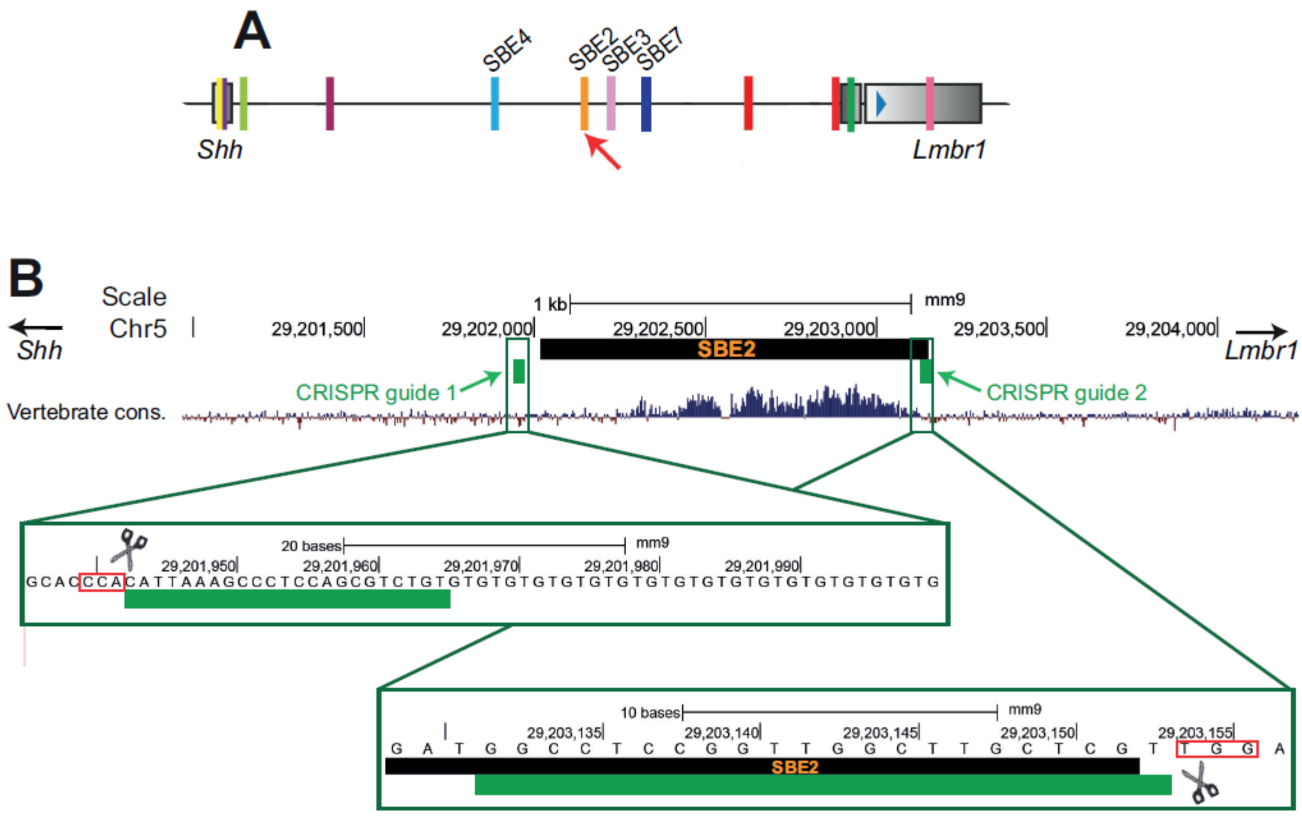


**Supplementary Figure 1. Overview of the region targeted via CRISPR for removal of SBE2 activity.** (A) Depiction of the *Shh* regulatory region. Coloured blocks represent the known *Shh* enhancers with the relevant brain enhancers labelled. The red arrow points to the location of the targeted SBE2 enhancer. (B) UCSC publicly available track of the vertebrate conservation for SBE2 and the neighbouring genomic region (https://genome.ucsc.edu/cgi-bin/hgTrackUi?db=hg19&g=cons100way). CRISPR guides are represented by the green blocks. A zoomed in view of each CRISPR guide and the targeted sequence is presented in the green boxes. The pam sites are demarcated by the red boxes.

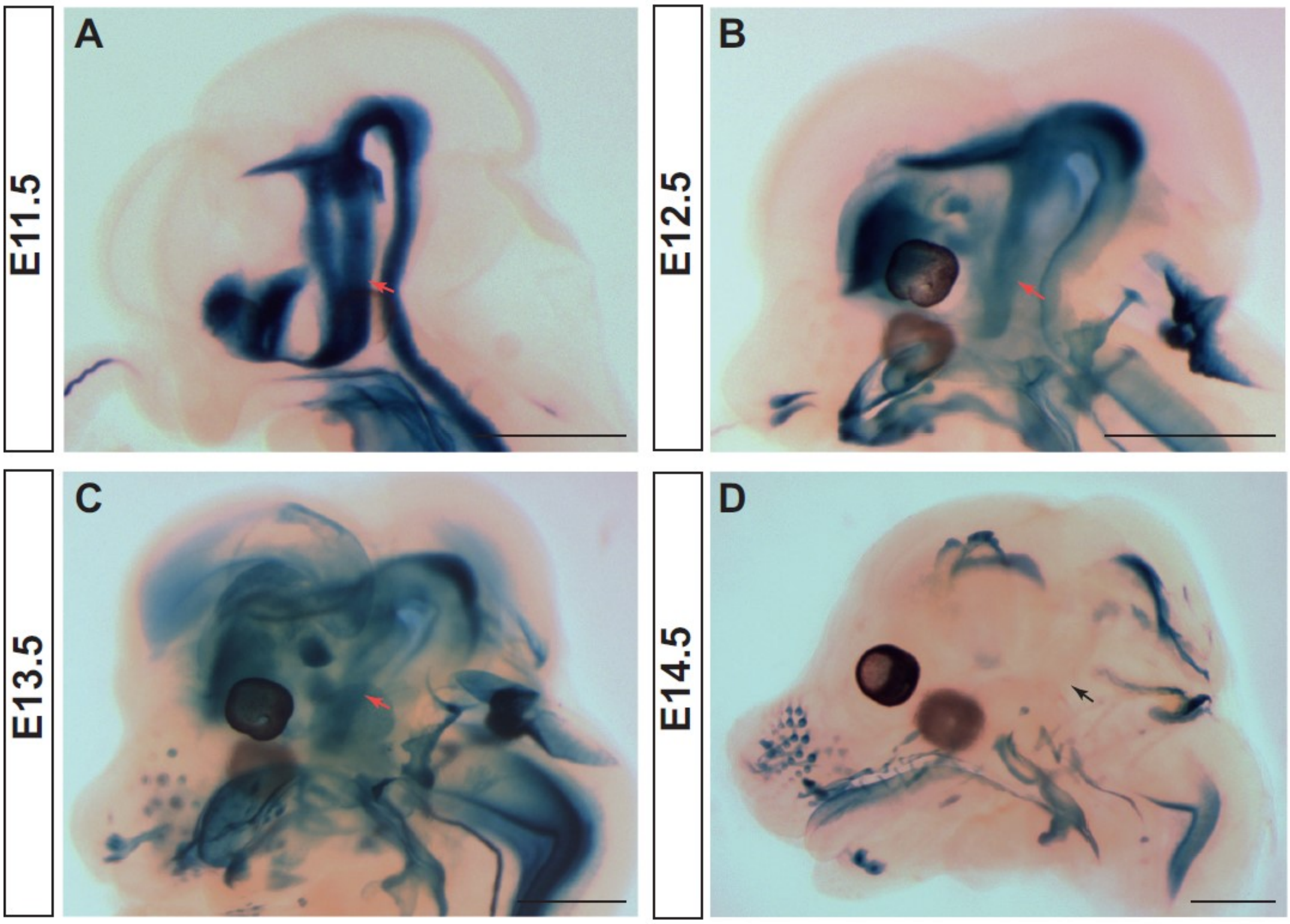


**Supplementary Figure 2. *Shh* expression is present within the VH up to E13.5.** (A-D) LacZ reporter gene expression in E11.5 (A), E12.5 (B), E13.5 (C) and E14.5 (D) embryonic heads of SBLacZ526 embryos representing regions of *Shh* expression. Red arrows point to VH expression, while the black arrow points to the VH region where no expression is seen at E14.5. (n=3)

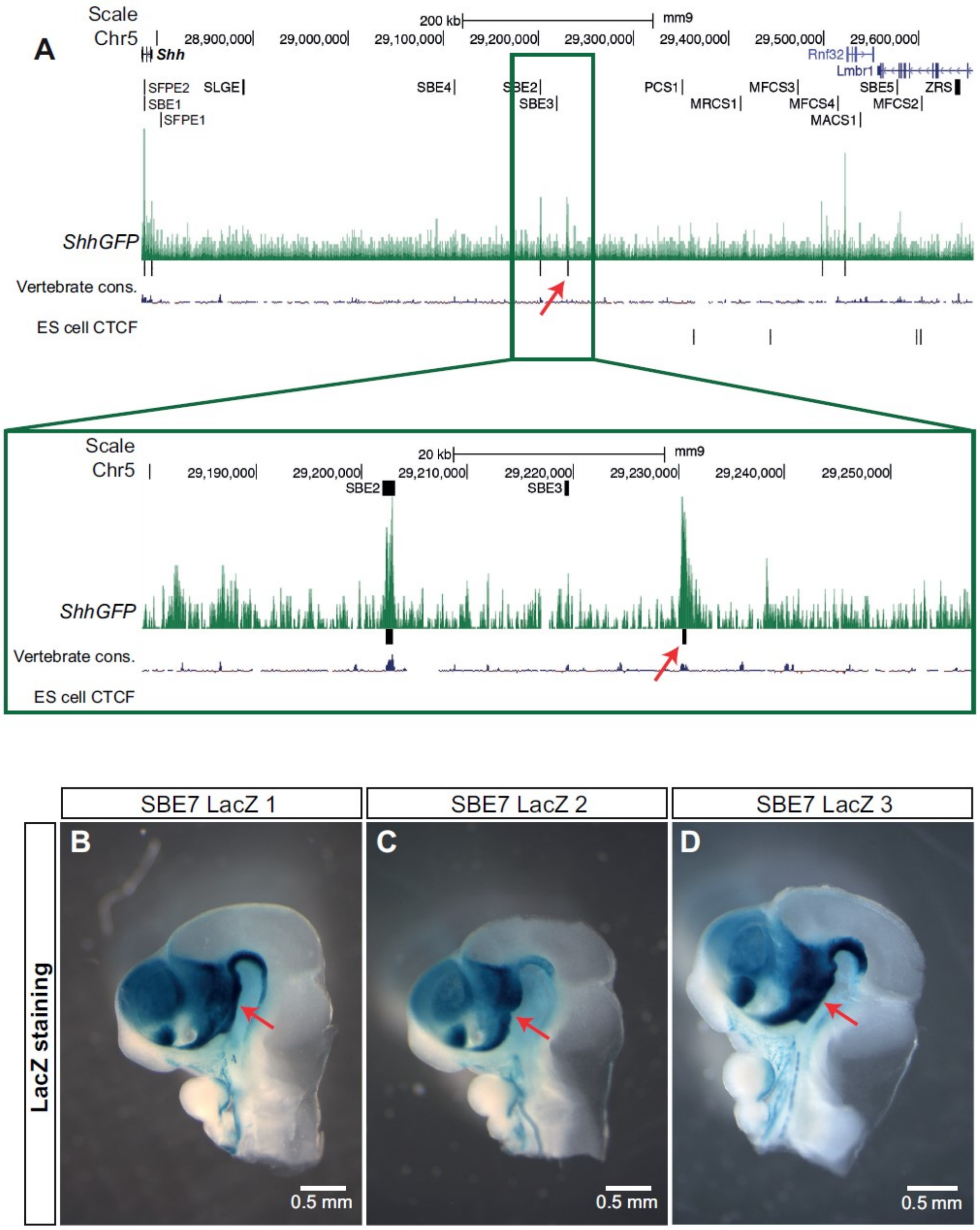


**Supplementary Figure 3. SBE7 is likely to drive residual *Shh* expression seen in the rostral diencephalon upon removal of SBE2 activity.** (A) UCSC track of ATAC-seq data from GFP positive cells from the entire head of *ShhGFP* embryos. The publicly available vertebrate conservation and sorted from BRUCE-4 ES cell peak tracks are also presented as are the sites for the known conserved enhancers. The red arrows in A point to the peak detected in *Shh* expressing cells of the brain identified as SBE7. A zoomed in view of the detected peak is presented in the green box more closely depicting the conservation seen in this region. (B-D) LacZ expression in E10.5 day heads (bisected along the midline) from three transgenics is shown, where LacZ expression is under the control of SBE7 from (A). Red arrows (in B-D )point to the region of the rostral diencephalon where LacZ expression is detected. Scale bars are shown in the bottom right hand corner.

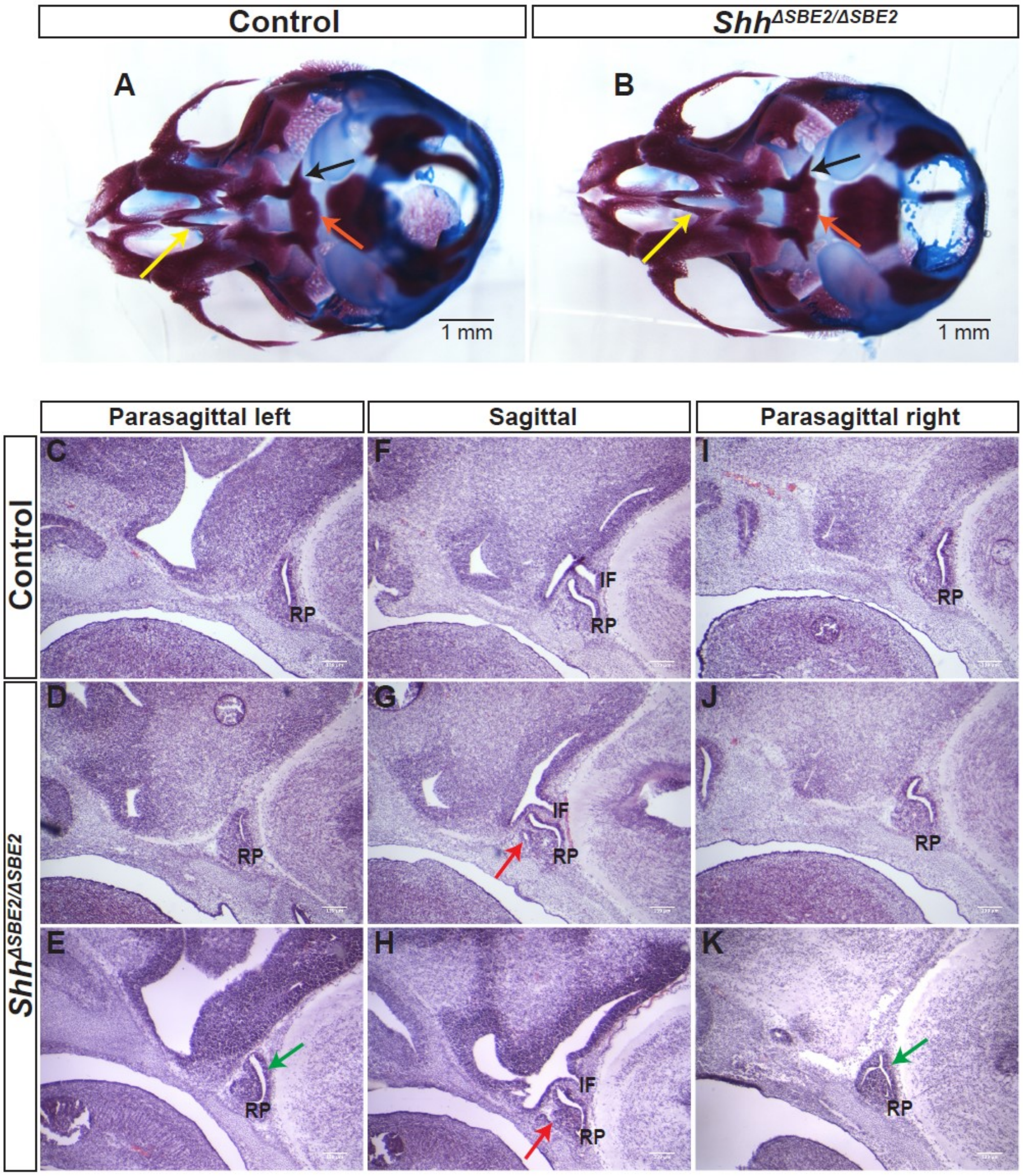


**Supplementary Figure 4. *Shh*$^{\Delta SBE2/\Delta SBE2}$ embryos lack deformities of the bones of the cranial vault and present mild lateral pituitary deformations.** (A-B) Dual staining highlighting chondrogenic (blue) and skeletal (red) craniofacial elements of control and *Shh*$^{\Delta SBE2/\Delta SBE2}$ E17.5 heads are shown. Orange arrows point to the basisphenoid bone, yellow arrows indicate the vomer bone and the black arrows point to the pterygoid bone. (C-K) H&E staining of E13.5 day left and right parasagittal and sagittal cryosections from control and two *Shh*$^{\Delta SBE2/\Delta SBE2}$ heads. Green arrows point to the mild lateral adenohypophyseal deformities seen in some *Shh*$^{\Delta SBE2/\Delta SBE2}$ mutant embryo. Red arrows point to the subtle midline adenohypophyseal deformities seen in *Shh*$^{\Delta SBE2/\Delta SBE2}$ embryos. RP- Rathke's pouch; IF- infundibulum. Scale bars are shown in the bottom right hand corner. (n=3)

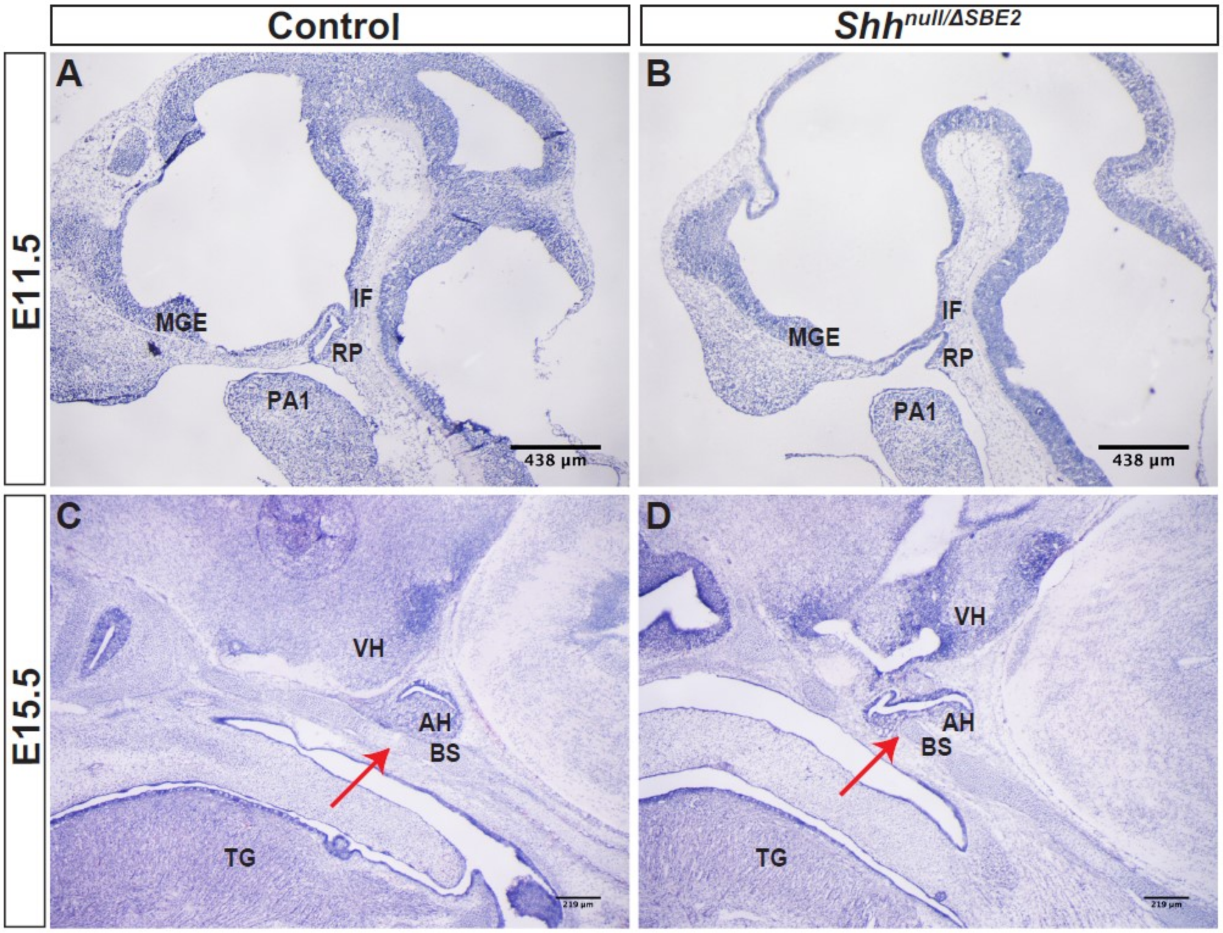


**Supplementary Figure 5.** ***Shh**$^{null/\Delta SBE2}$* **embryos present ventrally shifted pituitary lobes which impede basisphenoid bone formation.** (A-B) H&E stained control and *Shh*$^{null/\Delta SBE2}$ E11.5 head cryosections are shown. (C-D) H&E stained control and *Shh*$^{null/\Delta SBE2}$ E15.5 heads cryosections are depicted. Red arrows point to the region of the basisphenoid bone intercepted by the adenohypophysis in mutant embryos. RP- Rathke's pouch; IF- infundibulum; MGE- medial ganglionic eminence; PA1- pharyngeal arch 1; VH- ventral hypothalamus; BS- basisphenoid bone; TG- tongue; AH- adenohypophysis. Scale bars are shown in the bottom right hand corner. (n=3)

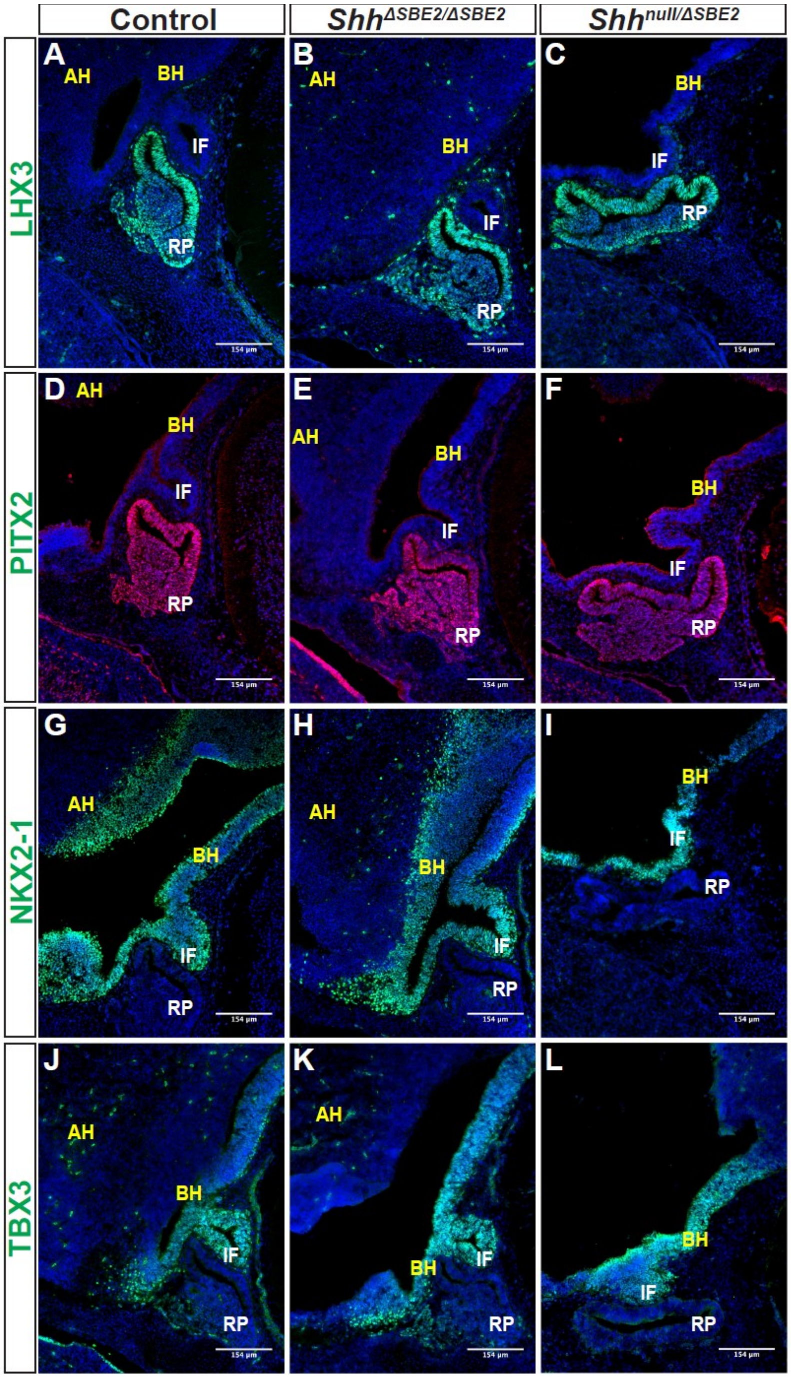


**Supplementary Figure 6. Patterning within the pituitary lobes is unaffected by loss of SBE2.** (A-C) Immunofluorescent staining for LHX3 in E13.5 day hypophyseal parasagittal cryosections of control, *Shh*$^{\Delta SBE2/\Delta SBE2}$ and *Shh*$^{null/\Delta SBE2}$ embryos. (D-F) Immunofluorescent staining for PITX2 in E13.5 day hypophyseal parasagittal cryosections of control, *Shh*$^{\Delta SBE2/\Delta SBE2}$ and *Shh*$^{null/\Delta SBE2}$ embryos. (G-I) Immunofluorescent staining for NKX2-1 in E13.5 day hypophyseal parasagittal cryosections of control, *Shh*$^{\Delta SBE2/\Delta SBE2}$ and *Shh*$^{null/\Delta SBE2}$ embryos. (J-L) Immunofluorescent staining for TBX3 in E13.5 day hypophyseal parasagittal cryosections of control, *Shh*$^{\Delta SBE2/\Delta SBE2}$ and *Shh*$^{null/\Delta SBE2}$ embryos. AH- alar hypothalamus; BH- basal hypothalamus; IF- infundibulum; RP- Rathke's pouch. Scale bars are shown in the bottom right hand corner. (n=3)

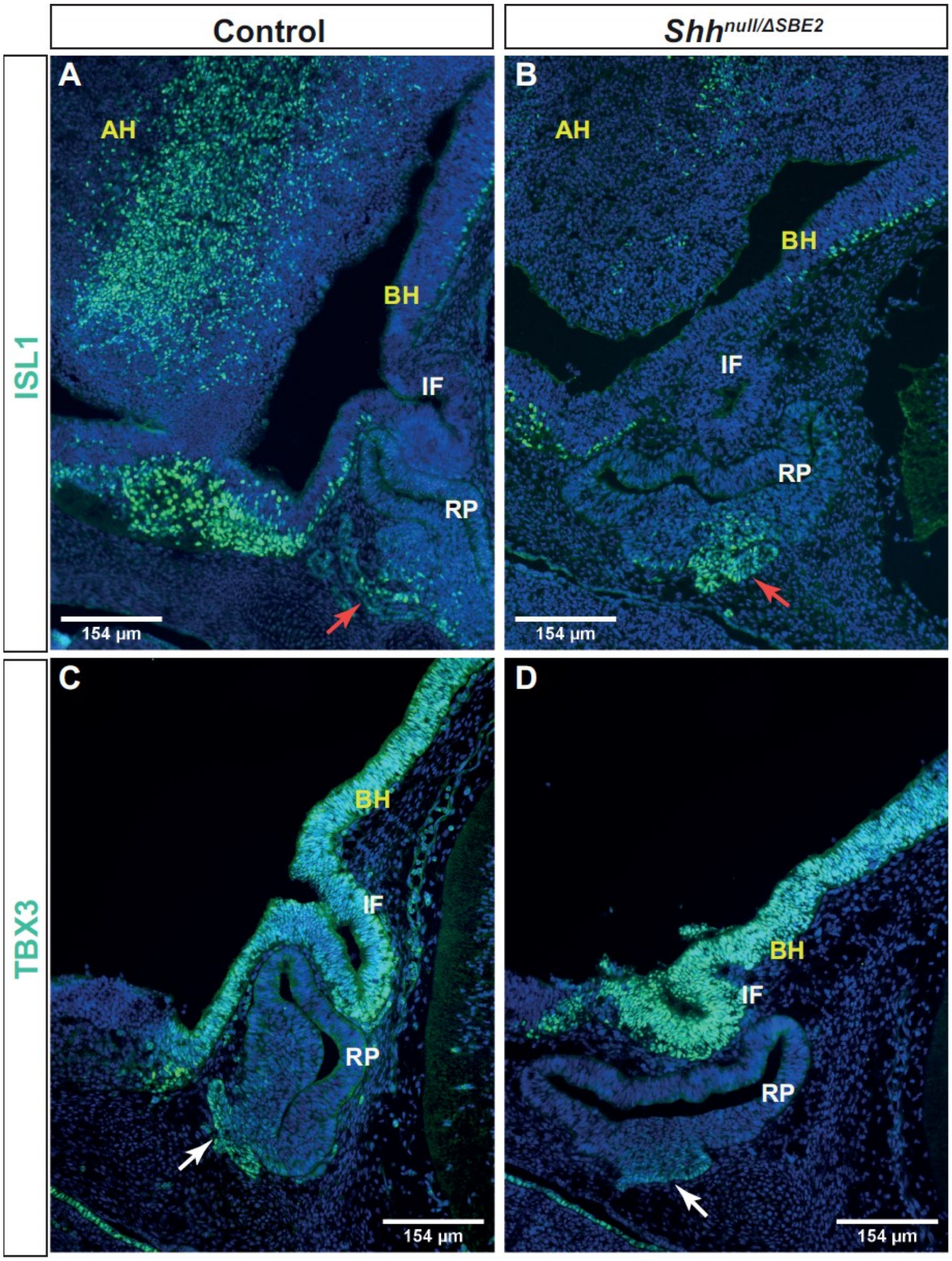


**Supplementary Figure 7. Loss of SBE2 does not lead to developmental delay of the adenohypophysis.** (A-B) Immunofluorescent staining for ISL1 in E12.5 day hypophyseal sagittal cryosections of control and *Shh*$^{null/\Delta SBE2}$ embryos. Red arrows point to the positive cell population in the ventral most portion of the adenohypophysis. (C-D) Immunofluorescent staining for TBX3 in E12.5 day hypophyseal parasagittal cryosections of control and *Shh*$^{null/\Delta SBE2}$ embryos. White arrows point to the positive staining in the ventral region of the adenohypophysis. BH- basal hypothalamus; IF- infundibulum; RP- Rathke's pouch. Scale bars are shown in the bottom left/right hand corner. (n=3)

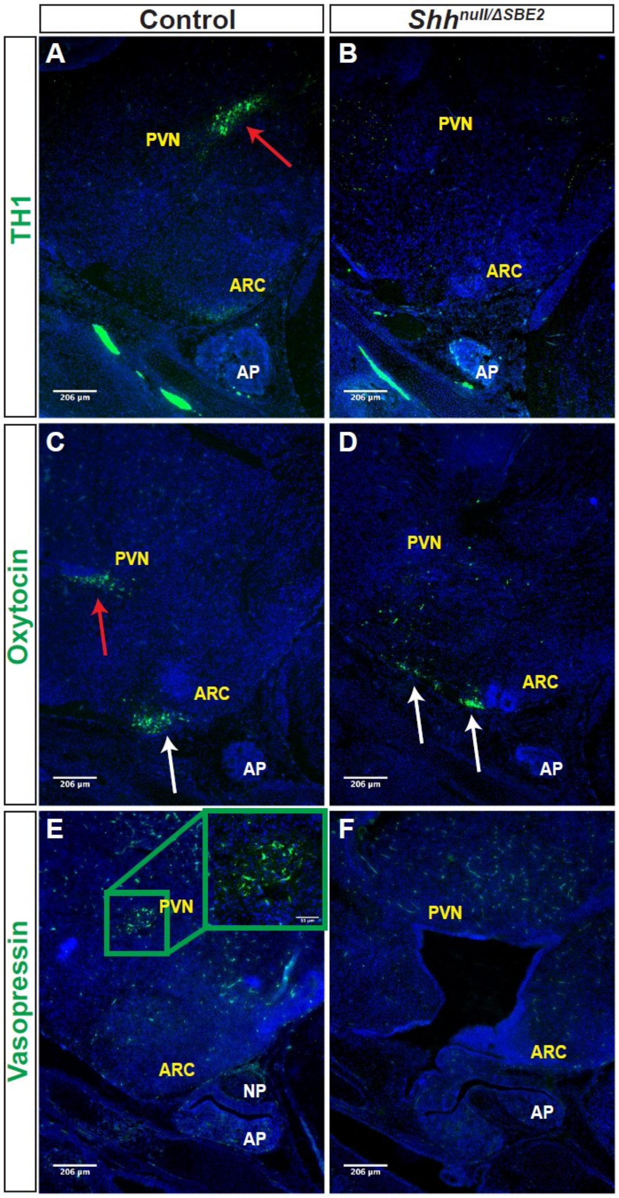


**Supplementary Figure 8. Neuronal populations which communicate with the hypophyseal lobes are affected in *Shh*$^{null/\Delta SBE2}$ embryos.** (A-B) Immunofluorescent staining for TH1 in E17.5 day VH parasagittal cryosections of control and *Shh*$^{null/\Delta SBE2}$ embryos. Red arrows point to the PVN TH1 neuronal population which is absent in *Shh*$^{null/\Delta SBE2}$ embryos. (C-D) Immunofluorescent staining for oxytocin in E17.5 day VH parasagittal cryosections of control and *Shh*$^{null/\Delta SBE2}$ embryos. Red arrows point to the PVN oxytocin neuronal population which is lost in *Shh*$^{null/\Delta SBE2}$ embryos. White arrows point to the ARC oxytocin neuronal population which is mislocalised in *Shh*$^{null/\Delta SBE2}$ embryos. (E-F) Immunofluorescent staining for vasopressin in E17.5 day VH parasagittal cryosections of control and *Shh*$^{null/\Delta SBE2}$ embryos. The green box presents the region of PVN expression of vasopressin which is absent in mutant embryos. A zoomed in view of the region of expression presented is seen in the upper right hand corner. NP- neurohypophysis; AP- adenohypophysis; PVN- paraventricular nucleus; ARC- arcuate nucleus. Scale bars are shown in the bottom left hand corner. (n=3)

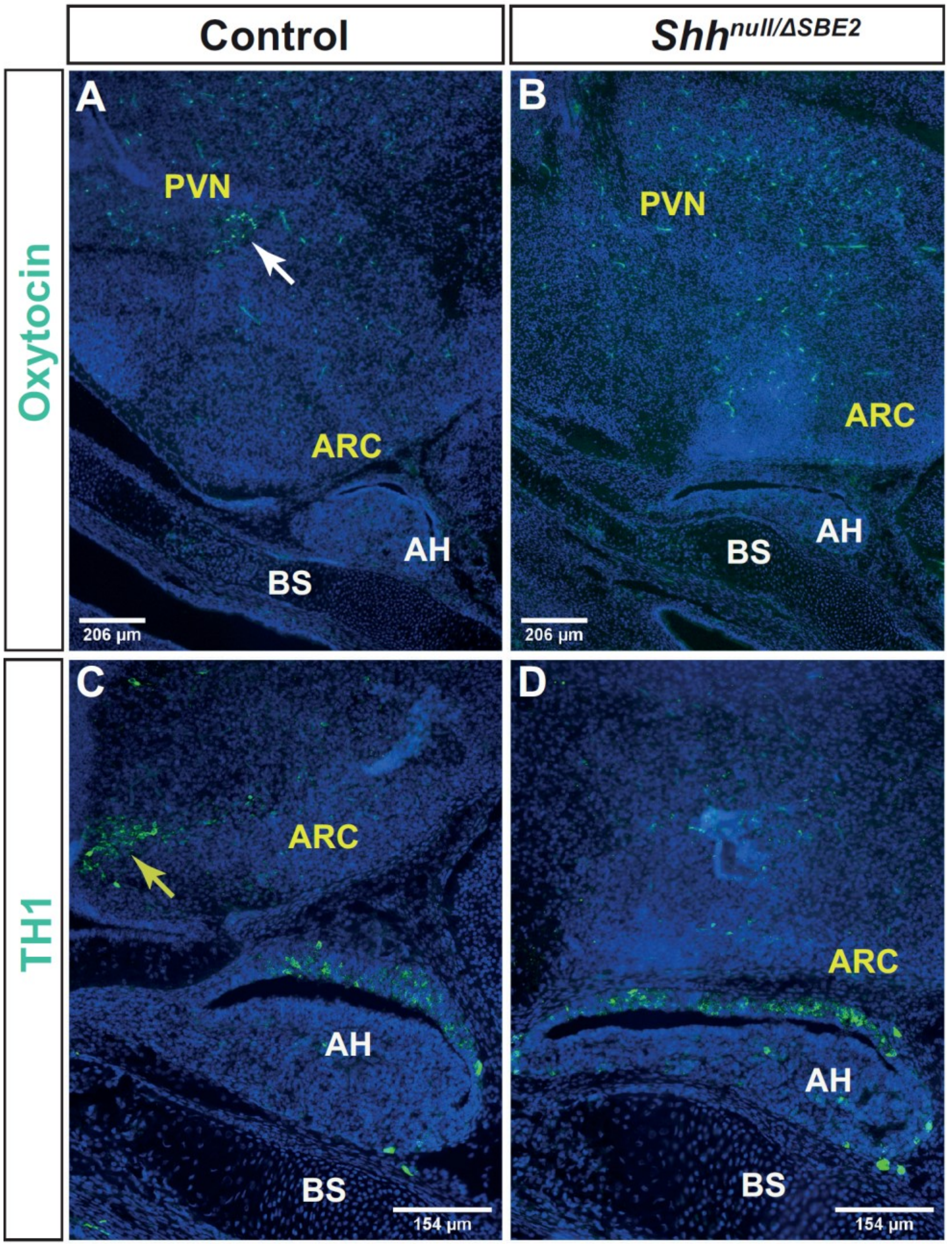


**Supplementary Figure 9. Loss of SBE2 does not cause a delay in hypothalamic neuronal specification.** (A-B) Immunofluorescent staining for oxytocin in E15.5 day VH parasagittal cryosections of control and $Shh^{null/\Delta SBE2}$ embryos. The white arrow points to the oxytocin population, lost in mutant embryos. (C-D) Immunofluorescent staining for TH1 in E15.5 day VH parasagittal cryosections of control and $Shh^{null/\Delta SBE2}$ embryos. The yellow arrow indicates the TH1 population, absent in mutant embryos. BS- basisphenoid; AH- adenohypophysis; PVN- paraventricular nucleus; ARC- arcuate nucleus. Scale bars are shown in the bottom left hand corner. (n=3)

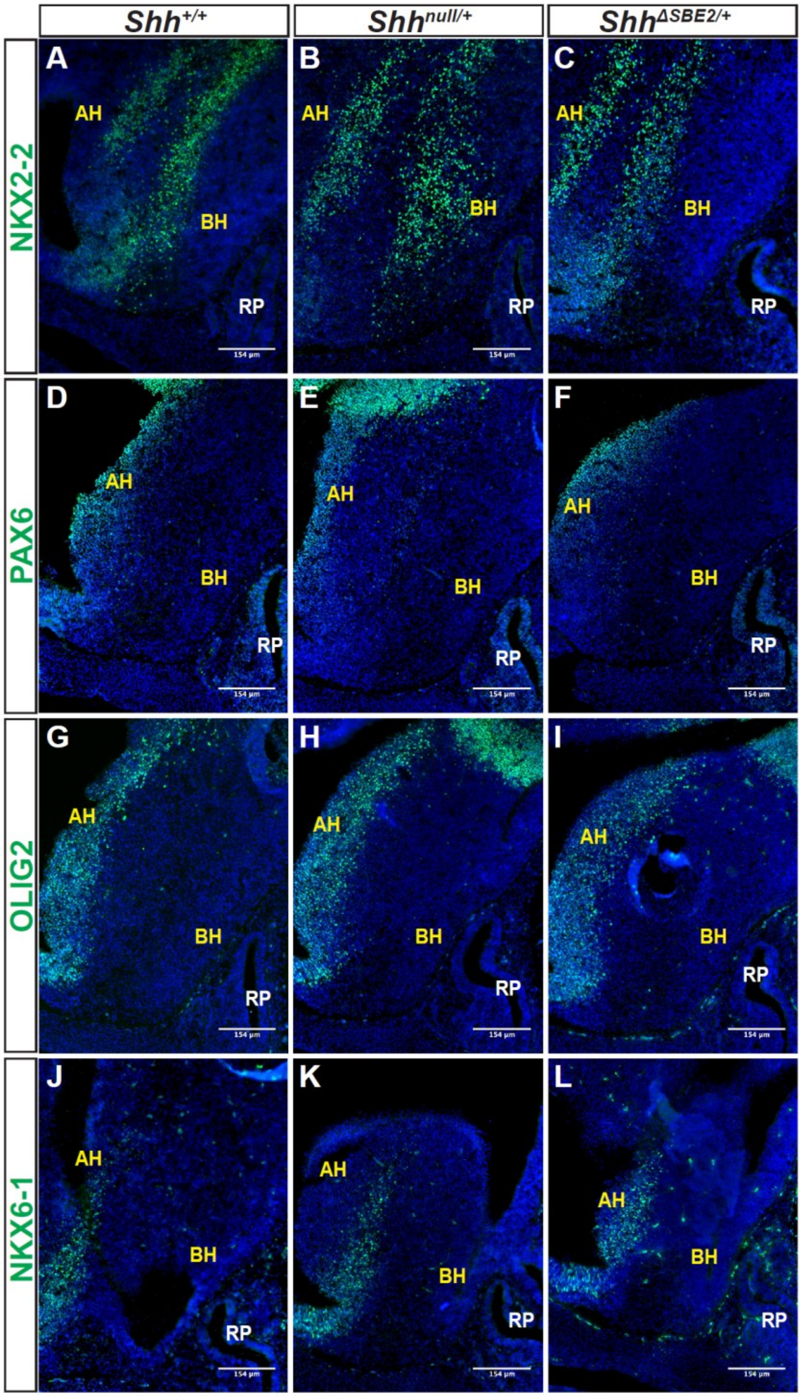


**Supplementary Figure 10. DV hypothalamic patterning is unaffected by heterozygous removal of SBE2 or *Shh* activity.** (A-C) Immunofluorescent staining for NKX2-2 in E13.5 day VH parasagittal cryosections of $Shh^{+/+}$, $Shh^{null/+}$ and $Shh^{\Delta SBE2/+}$ embryos. (D-F) Immunofluorescent staining for PAX6 in E13.5 day VH parasagittal cryosections of $Shh^{+/+}$, $Shh^{null/+}$ and $Shh^{\Delta SBE2/+}$ embryos. (G-I) Immunofluorescent staining for OLIG2 in E13.5 day VH parasagittal cryosections of $Shh^{+/+}$, $Shh^{null/+}$ and $Shh^{\Delta SBE2/+}$ embryos. (J-L) Immunofluorescent staining for NKX6-1 in E13.5 day VH parasagittal cryosections of $Shh^{+/+}$, $Shh^{null/+}$ and $Shh^{\Delta SBE2/+}$ embryos. AH- alar hypothalamus; BH- basal hypothalamus; RP- Rathke's pouch. Scale bars are shown in the bottom right hand corner. (n=3)

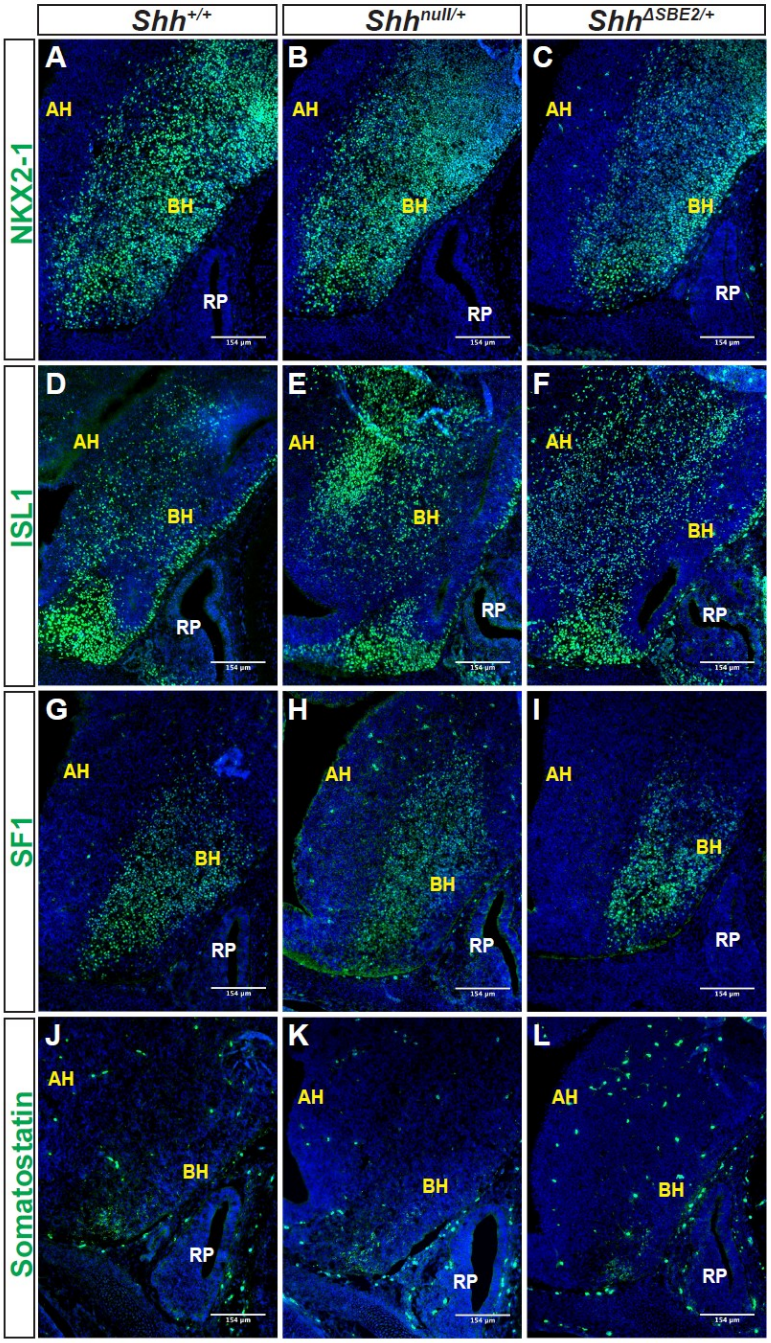


**Supplementary Figure 11. DV hypothalamic neuronal specification is unaffected by heterozygous removal of SBE2 or *Shh* activity.** (A-C) Immunofluorescent staining for NKX2-1 in E13.5 day VH parasagittal cryosections of $Shh^{+/+}$, $Shh^{null/+}$ and $Shh^{\Delta SBE2/+}$ embryos. (D-F) Immunofluorescent staining for ISL1 in E13.5 day VH parasagittal cryosections of $Shh^{+/+}$, $Shh^{null/+}$ and $Shh^{\Delta SBE2/+}$ embryos. (G-I) Immunofluorescent staining for SF1 in E13.5 day VH parasagittal cryosections of $Shh^{+/+}$, $Shh^{null/+}$ and $Shh^{\Delta SBE2/+}$ embryos. (J-L) Immunofluorescent staining for somatostatin in E13.5 day VH parasagittal cryosections of $Shh^{+/+}$, $Shh^{null/+}$ and $Shh^{\Delta SBE2/+}$ embryos. AH- alar hypothalamus; BH- basal hypothalamus; RP- Rathke's pouch. Scale bars are shown in the bottom right hand corner. (n=3)
